# Do congruent lip movements facilitate speech processing in a dynamic audiovisual multi-talker scenario? An ERP study with older and younger adults

**DOI:** 10.1101/2020.11.06.370841

**Authors:** Alexandra Begau, Laura-Isabelle Klatt, Edmund Wascher, Daniel Schneider, Stephan Getzmann

**Affiliations:** Leibniz Research Centre for Working Environment and Human Factors, TU Dortmund, Germany

**Keywords:** audiovisual integration, multi-talker speech perception, aging, attention, event-related potentials, multisensory processing

## Abstract

In natural conversations, visible mouth and lip movements play an important role in speech comprehension. There is evidence that visual speech information improves speech comprehension, especially for older adults and under difficult listening conditions. However, the neurocognitive basis is still poorly understood. The present EEG experiment investigated the benefits of audiovisual speech in a dynamic cocktail-party scenario with 22 (aged 20 to 34 years) younger and 20 (aged 55 to 74 years) older participants. We presented three simultaneously talking faces with a varying amount of visual speech input (still faces, visually unspecific and audiovisually congruent). In a two-alternative forced-choice task, participants had to discriminate target words (“yes” or “no”) among two distractors (one-digit number words). In half of the experimental blocks, the target was always presented from a central position, in the other half, occasional switches to a lateral position could occur. We investigated behavioral and electrophysiological modulations due to age, location switches and the content of visual information, analyzing response times and accuracy as well as the P1, N1, P2, N2 event-related potentials (ERPs) and the contingent negative variation (CNV) in the EEG. We found that audiovisually congruent speech information improved performance and modulated ERP amplitudes in both age groups, suggesting enhanced preparation and integration of the subsequent auditory input. In the older group, larger amplitude measures were found in early phases of processing (P1-N1). Here, amplitude measures were reduced in response to audiovisually congruent stimuli. In later processing phases (P2-N2) we found decreased amplitude measures in the older group, while an amplitude reduction for audiovisually congruent compared to visually unspecific stimuli was still observable. However, these benefits were only observed as long as no location switches occurred, leading to enhanced amplitude measures in later processing phases (P2-N2). To conclude, meaningful visual information in a multi-talker setting, when presented from the expected location, is shown to be beneficial for both younger and older adults.

## 1. Introduction

Imagine meeting for lunch with a group of people. You ask your colleague from across the table about their weekend, but you can barely understand what they are saying because next to you, other people are also chatting. It’s long known that verbal conversation in loud and noisy environments and in the presence of concurrent talkers is hard work for our auditory system (Alain, McDonald, Ostroff, & Schneider, 2004). In such “cocktail-party” situations (Cherry, 1953), extracting relevant speech content from a multitude of speech streams mainly demands two cognitive processes: Firstly, the segregation of target speech against the irrelevant speech streams and secondly, the streaming of speech or the connection of speech elements across time (Bronkhorst, 2015; Hill & Miller, 2010). Such a pre-attentive semantic processing of speech, also called grouping, is required to trigger attentive processes and to guide selective auditory attention (Bronkhorst, 2015; Carlile, 2014). Attentional demands are highly modulated by the listening situation itself. For example, “homing in” auditory attention onto a talker of interest takes some time, and therefore speech comprehension is enhanced when the spatial location of the perceived target remains the same (Best, Ozmeral, Kopco, & Shinn-Cunningham, 2008). In everyday life, however, conversational situations are often highly dynamic and changing speakers pose a particular challenge for the auditory system as switches in talker locations require a continuous re-focusing of attention (Koch, Lawo, Fels, & Vorländer, 2011; Lawo, Fels, Oberem, & Koch, 2014; Lin & Carlile, 2015).

Nevertheless, speech has several advantageous characteristics that makes it distinguishable from other acoustic signals and therefore especially suitable for situations with multiple talkers. It proves resistant to background masking, making a single speech stream stand out amongst the noise. The high redundancy of speech provides a “safety net”, enabling the listener to fill in the gap where information is missing due to noise or background babble (for review, see Bronkhorst, 2015). Expectations and prior knowledge about the speech content play an important role in these situations. Also, the familiarity of a talker’s voice (e.g. Johnsrude et al., 2013) and the anticipation of a target talker’s location among concurrent talkers (Kidd, Arbogast, Mason, & Gallun, 2005) are used to filter out relevant information out of a mixture of auditory input.

A further important source of redundancy of speech, which is often not taken into account in studies on speech comprehension, is visual speech information. In natural speech, the expectations concerning the auditory input are modulated by complementary visual information (Lindström, 2012; van Wassenhove, Grant, & Poeppel, 2005), enhancing both audiovisual processing (Besle, Fort, Delpuech, & Giard, 2004; for review see Campbell, 2008; Tye-Murray, Spehar, Myerson, Hale, & Sommers, 2016; Winneke & Phillips, 2011) and speech intelligibility (Brault, Gilbert, Lansing, McCarley, & Kramer, 2010). For audiovisual speech integration to be successful, the typically preceding visual stimulus needs to be a valid predictor of the auditory input (Stekelenburg & Vroomen, 2007; van Wassenhove et al., 2005), in line with the analysis-by-synthesis model by van Wassenhove et al. (2005). In order to perceive auditory and visual speech as one coherent audiovisual event, it has been suggested that a general binding process of both modalities occurs first, followed by temporal, spatial and finally phonemic integration of auditory and visual speech (Baart, Stekelenburg, & Vroomen, 2014; Ganesh, Berthommier, Vilain, Sato, & Schwartz, 2014).

Thus, integration of both modalities happens early in the process of speech perception, even pre-phonemically, which leads to a better performance (Besle et al., 2004; Campbell, 2008; van Wassenhove et al., 2005) possibly due to freed cognitive resources enabling a more precise interpretation of the input (Campbell, 2008). Benefits of audiovisual speech-processing are therefore especially pronounced under difficult listening conditions (Grant & Seitz, 2000; Heald & Nusbaum, 2014; Schwartz, Berthommier, & Savariaux, 2004).

Audiovisual benefits should also be evident when individual prerequisites for undisturbed speech comprehension are not fulfilled, for example, in people with hearing disorders or at an advanced age. Especially age-related declines in hearing abilities and in the processing of speech can be regarded as a prototypical example of how audiovisual input could enhance speech comprehension. Decline occurs on both the sensory and cognitive level: Hearing loss (Presbycusis) and neuronal changes, resulting in a deteriorated input of sensory information (see Peelle & Wingfield, 2016), declines in general attentional mechanisms, working memory, and processing speed (e.g. Burke & Shafto, 2015; Kropotov, Ponomarev, Tereshchenko, Müller, & Jäncke, 2016), as well as a reduced auditory spatial attention (Dai, Best, & Shinn-Cunningham, 2018) contribute to age-related hearing issues. As a consequence, older adults are usually less able to perform well in attentional highly demanding situations such as “cocktail-party” environments, where they experience difficulties with regulating their attentional focus toward the relevant stream of auditory information (Passow et al., 2012). When irregular spatial shifts of the target occur, older adults are even more affected by distractors than younger people (Volosin, Gaál, & Horváth, 2017) and need longer phases of (re-)orientation toward the relevant stimulus (Correa-Jaraba, Cid-Fernández, Lindín, & Díaz, 2016; Getzmann, Falkenstein, & Wascher, 2015).

On the other hand, older people make use of several compensation strategies like the enhancement of preparatory and orienting processes (Kropotov et al., 2016) and stronger top-down regulation of the attentional focus, keeping the behavioral performance relatively stable (Alain et al., 2004). Accordingly, several studies showed an age-related posterior-to-anterior shift in neuronal activation, suggesting compensatory mechanisms based on enhanced engagement of fronto-central brain areas (Davis, Dennis, Daselaar, Fleck, & Cabeza, 2008; Kropotov et al., 2016). Moreover, audiovisual integration is still intact with age, even though visual acuity (see Owsley, 2011) as well as the processing of visual speech, that is, the ability to lipread, appears to be diminished in older adults (Sommers, Tye-Murray, & Spehar, 2005; Tye-Murray et al., 2016; Winneke & Phillips, 2011). Still, the audiovisual performance is similar, indicating that older adults might profit even more from multimodal speech than younger adults (Winneke & Phillips, 2011). Moreover, older adults rely more on the visual information, especially when the auditory information is ambiguous (Cienkowski & Carney, 2002; Sekiyama, Soshi, & Sakamoto, 2014). At the same time, the quality of the visual content seems to gain importance. While younger adults profit from any meaningful visual content, older adult’s performance is supported by high quality visual input only (Tye-Murray, Spehar, Myerson, Sommers, & Hale, 2011).

Taken together, previous research has shown that audiovisual speech perception is important, especially under difficult listening conditions and for older people. However, studies exploring audiovisual speech integration in younger and older adults in dynamic “cocktail-party” situations are scarce and have mainly focused on behavioral measures. In the present study, we investigated the underlying neuro-cognitive underpinnings of audiovisual multi-talker speech processing, using EEG-related measures in combination with a dynamic speech perception paradigm. We focused our analysis on a number of event-related potentials (ERPs) that have already been studied in relation to both audiovisual integration and multi-talker scenarios. In general, auditory ERPs like P1, N1 and P2 have been linked to sensory detection and the cognitive processing of sound (for a review, see Alain & Tremblay, 2007). The first component, the P1, a rather small positive deflection in central scalp regions is modulated by physical characteristics of the auditory stimulus (Alain & Tremblay, 2007) and is thought to reflect early processing of sensory information (Baart, 2016). In the presence of additional visual information, the P1 is usually reduced in amplitude (Stothart & Kazanina, 2016; Winneke & Phillips, 2011), possibly reflecting cross-modal sensory gating, that is, a more effective pre-attentive filtering of sensory information (Lebib, Papo, de Bode, & Baudonnière, 2003). Accordingly, larger P1 amplitudes can be found, when easily detectable incongruent audiovisual input is presented, indicating an early mechanism to detect non-redundant audiovisual information. Larger amplitudes have also been found in older compared to younger adults, which has been linked to an age-related deficit in inhibition or gating (Friedman, 2011; Getzmann et al., 2015; Stothart & Kazanina, 2016; Winneke & Phillips, 2011).

Following the P1, there is a negative and a subsequent positive deflection called the N1-P2 complex (Alain & Tremblay, 2007). Baart (2016) investigated the influence of visual speech on N1 and P2 in a meta-analysis and found an overall suppression and speeding-up of these components throughout different studies. Van Wassenhove and colleagues (2005) argued within their analysis-by-synthesis model that auditory and visual information interact as early as at the N1-latency. The predictive value of the visual information for the auditory input seems to influence the degree of temporal facilitation in both components. That is, the more salient and predictable the visual input is, the stronger is the facilitation of the auditory processing. Accordingly, higher attentional demands seem to disrupt audiovisual integration early during processing, resulting in a decrement of the latency reduction of both components (Alsius, Möttönen, Sams, Soto-Faraco, & Tiippana, 2014). Importantly, both components also differ with respect to content-dependency. While the N1 is simply driven by anticipation of the auditory speech input due to the preceding visual input, a P2 modulation can only be observed, when the visual input reliably predicts the auditory input and speech specific binding can occur (Ganesh et al., 2014; Stekelenburg & Vroomen, 2007). It is therefore suggested, that the N1 reflects the integration of spatial and temporal audiovisual properties, while the P2 reflects the subsequent integration of phonetic properties (Baart et al., 2014; Klucharev, Möttönen, & Sams, 2003).

N1 is also thought to reflect the extent of distraction when a switch in talker location occurs. Volosin and colleagues (2017) found similar modulations in younger and older adults, arguing that older adults needed no extra time to recover from distraction. Contrarily, a study investigating distraction in a multi-talker scenario found an age-related slowing of both N1 and P2, as well as a P2 amplitude reduction with a present distractor only in younger adults (Getzmann et al., 2015). Taken together, as reviewed by Friedmann (2011), age-related modulations of N1 amplitudes are somewhat ambiguous throughout literature. Similarly, the P2 in relation to aging processes is not well assessed, although the literature also indicated an age-related enhancement and slowing of the component (Friedman, 2011).

Following the P1-N1-P2 complex, a later component, the N2 can be observed that has been linked to attentional control processes and conscious sensory discrimination (Friedman, 2011; Ritter, Simson, Vaughan, & Friedman, 1979). Processes contributing to a N2 modulation in both multimodal and multi-talker scenarios have yet to be unraveled. To date, the literature suggests a number of scarce or ambiguous findings. A larger N2 amplitude is correlated with better performance and faster reaction times in audiovisual processing (Lindström, 2012). In regard to age-related modulations of the N2 amplitude, both increases and decreases in amplitudes and consistent increases in latencies have been observed (for a review see Friedman, 2011). Several more recent studies showed a decline or absence of N2 amplitudes in older adults, in both audiovisual (Stothart & Kazanina, 2016) and multi-talker scenarios (Getzmann et al., 2015; Getzmann & Wascher, 2016), supporting the notion of an age-related inhibitory deficit. Stothart and Kazanina (2016) suggested that congruent visual information helps the elderly to process auditory input, while processing incongruent audiovisual information comes with greater conflict costs. Getzmann and Wascher (2016) found a decrease in N2 amplitude in older adults only when unexpected irrelevant information was presented, arguing that due to age, the control over irrelevant information is weakened.

In addition to these ERPs related to the onset of speech stimuli, a pronounced negative deflection immediately preceding the onset of auditory speech can be observed, which has been related to the Contingent Negative Variation (CNV, Walter, Cooper, Aldridge, McCallum, & Winter, 1964). The CNV reflects preparatory processes in expectation of an upcoming, task-relevant event. In a previous multi-talker speech perception study, for example, the CNV was reduced in older participants, which was interpreted as an age-related decline in preparatory activity of an upcoming auditory event (Getzmann, Golob, & Wascher, 2016). In the present literature concerning audiovisual processing, anticipatory slow waves such as the CNV are known, but usually not further investigated (as seen in Alsius et al., 2014; Klucharev et al., 2003). In fact, it is rather discussed as problematic for the interpretation of later ERP differences in terms of cross-modal interaction (Teder-Salejarvi et al., 2002).

To summarize, audiovisual speech perception involves a number of cognitive mechanisms resulting in an effective processing of both modalities and in several advantages compared to auditory-only speech. Here, we investigated the potential benefit of audiovisual speech information in cocktail-party scenarios, in which the listener is confronted with multiple simultaneously talking persons and occasional changes in talker locations.

Specifically, three talkers simultaneously uttered short words, one containing the target information and two containing concurrent speech input. We varied the information content of visual speech information (i.e., audiovisually congruent speech input, visually unspecific speech input, no speech input) in order to assess its influence on speech comprehension. In addition, we employed two different listening scenarios, a *static* setting, in which the target talker location always remained the same, and a *dynamic* setting, in which the target talker occasionally switched position with the concurrent talkers. The *dynamic* setting was used to explore the influence of changes in talker locations on the processing of audiovisual speech information. We analyzed both behavioral performance and electrophysiological measures of younger and older adults, expecting to see audiovisual facilitation in form of a faster and more accurate detection of the target information with congruent visual speech input, relative to visually unspecific or no visual input. We hypothesized older adults to show a worsened performance, that is, slower responses and lower accuracy, compared to younger adults. On the electrophysiological level, we expected reduced P1 amplitudes, when audiovisual congruent information is presented as well as modulations in N1 and P2 amplitudes. Here, we expected audiovisual congruency to be important only for the modulation of P2, but not for N1 amplitudes. In case of the N2, we expected larger amplitudes with audiovisually congruent information. Modulations should differ across age-groups. Even though the literature presents ambiguous results, we expected amplitude enhancements in the early P1-N1-P2 complex and a decrease or absence of N2 amplitudes in the older group due to an age-related inhibitory deficit. In addition to these ERPs triggered by the onset of the acoustic speech stimulus, preparatory activity reflected by a slow-wave potential such as the CNV is likely to occur, especially with audiovisual congruent information. As stated by Baart (2016), similar studies on audiovisual processing oftentimes focus on single component measures, however the methodological assessment is not consistent. In order to account for possible distortions in the single ERPs due to a preceding CNV, we focused on amplitude to amplitude measures (as in several previous studies, e.g. Kokinous, Kotz, Tavano, & Schröger, 2015; Simon & Wallace, 2018). Hence, processing was analyzed in three stages: early, reflected by P1 to N1 measures, followed by N1 to P2, and finally late processing, reflected by P2 to N2. In terms of these three amplitude to amplitude measures, we expect to observe an age-related enhancement in P1-N1 and N1-P2 as well as a decrease in P2-N2. We expect audiovisual congruent information to reduce the P1-N1, but also to modulate the following measures N1-P2 and P2-N2. Finally, with regard to the *dynamic* scenario, we expected a reduced benefit of visual speech input when a change in target talker location occurred, given that the visual information presented at the expected target position cannot be used for integration with the auditory target speech information.

## 2. Method

### 2.1 Participants

50 participants were invited to participate in the study. Eight participants had to be excluded due to severe hearing problems (*n* = 4), visual problems (*n* = 1), technical issues (*n* = 2), and missing compliance to finish the experiment (*n* = 1). Thus, data of 42 participants could be analyzed, 22 of them were aged 20 to 34 years (*M_young_* = 25.45, *SE_young_* = 0.70; 12 female) and 20 were aged 55 to 70 years (*M_old_* = 64.50, *SE_old_* = 0.98; 11 female). All participants reported to be right-handed and mentally fit without any history of neurological diseases or any psychopharmacological medication. Before starting any testing, each participant gave written informed consent. They received a payment of 10€ per hour at the end of the experiment. The experimental procedure was in accordance with the Declaration of Helsinki, approved by the Ethical Committee of the Leibniz Research Centre for Working Environment and Human Factors, Dortmund, Germany.

### 2.2 Sensory and cognitive abilities

Hearing performance was assessed with a pure-tone audiometry (Oscilla USB 330; Inmedico, Lystrup, Denmark) at eleven pure-tone frequencies in-between 125–8000 Hz. Participants mostly achieved a hearing threshold ≤ 30 dB below 4000 Hz, even though mild to moderate presbycusis was observed in the older group. Mild hearing loss was observed in several participants. At 2000 Hz, thresholds were ≤ 35 dB for two participants. At 3000 Hz, we observed thresholds ≤ 35 dB (*n* = 2), ≤ 40 dB (*n* = 2) and ≤ 45 dB (*n* = 1). Thus, the older group had significantly lower hearing levels, that is a lower mean across both ears for frequencies 125 to 4000 Hz, than the younger group, *M_old_* = 16.81, *SE_old_* = 0.93*, M_young_* = 9.03, *SE_young_* = 0.60, *t*(39) = 7.07, *p* < .001, *g* = 2.17. Given the relatively loud stimulus presentation (75 dB), we considered this still acceptable.

Visual acuity was measured using *Landolt C optotypes* in 1.5 m distance and was at least 0.75 for both eyes in all participants, while a value of 1.0 is considered “normal” visual acuity (ISO 8596:2017(E)). Contrast sensitivity was assessed using the *Pelli-Robson Contrast Sensitivity Chart* in 1.0m distance (Pelli, Robson, & Wilkins, 1988). A score of 1.65 is recommended as lowest contrast sensitivity measure (Mäntyjärvi & Laitinen, 2001) and was achieved by all but one (older female) participant with a score of 1.5. Between-group comparisons revealed lower scores in visual acuity (*V* = 77.5, *p* <.001, *r* = .58) and contrast sensitivity (*V* = 68, *p* <.001, *r* = .65) in older compared to younger participants.

Cognitive functions were assessed using the *Montreal Cognitive Assessment* (MoCA; Nasreddine et al., 2005), a screening tool for Mild Cognitive Impairment. Older participants scored overall lower than younger ones (*M_old_* = 26.40, *SE_old_* = 0.62; *M_young_* = 28.68, *SE_young_* =0.29; *V* = 90.5, *p* < .001, *r* = .51), while all participants met the recommended cut-off criterion of 17 points as an indication for dementia (Carson, Leach, & Murphy, 2018). Working memory capacity was measured using the Digit Span subtest from the German Adaptation of the *Wechsler Adult Intelligence Scale* (Hamburg-Wechsler Intelligenztest für Erwachsene – Revision 1991; HAWIE-R, Tewes, 1991). Younger participants performed better than the older ones in forward recall (*M_young_* = 9.59 digits, *SE_young_* = 0.41, *M_old_* = 6.90, *SE_old_* = 0.46; *t*(39) = −4.37, *p* < .001, *g* = −1.34) and in backward recall (*M_young_* = 8.54 digits, *SE_young_* = 0.39, *M_old_* = 6.84, *SE_old_* = 0.45; *t*(39) = −2.72*, p* = .010, *g* = −0.84).

### 2.3 Materials and stimuli

Participants were seated in a dimly lit, sound attenuated room (5.0 × 3.3 × 2.4 m³). Attenuation was achieved through pyramid-shaped foam panels on ceiling and walls and a woolen carpet on the floor, resulting in a background noise level below 20 dB (A). For audiovisual stimulus presentation, we used three 12” vertically aligned monitors with a 1080 by 1920-pixel resolution and a 50 Hz refresh rate (Beetronics, Düsseldorf, Germany). The auditory signal was presented at about 75 dB SPL using three full range loudspeakers (SC 55.9 - 8 Ohm; Visaton, Haan, Germany), each mounted underneath a monitor. To achieve a free-field listening scenario, monitors and loudspeakers were installed on a horizontal array at −15° (left), 0° (center) and +15° (right) azimuth at 1.5m distance in approximate head height (lower frame at 1.00 m (loudspeaker) and 1.12 m (monitor), respectively, figure 1A).

**Figure 1.**
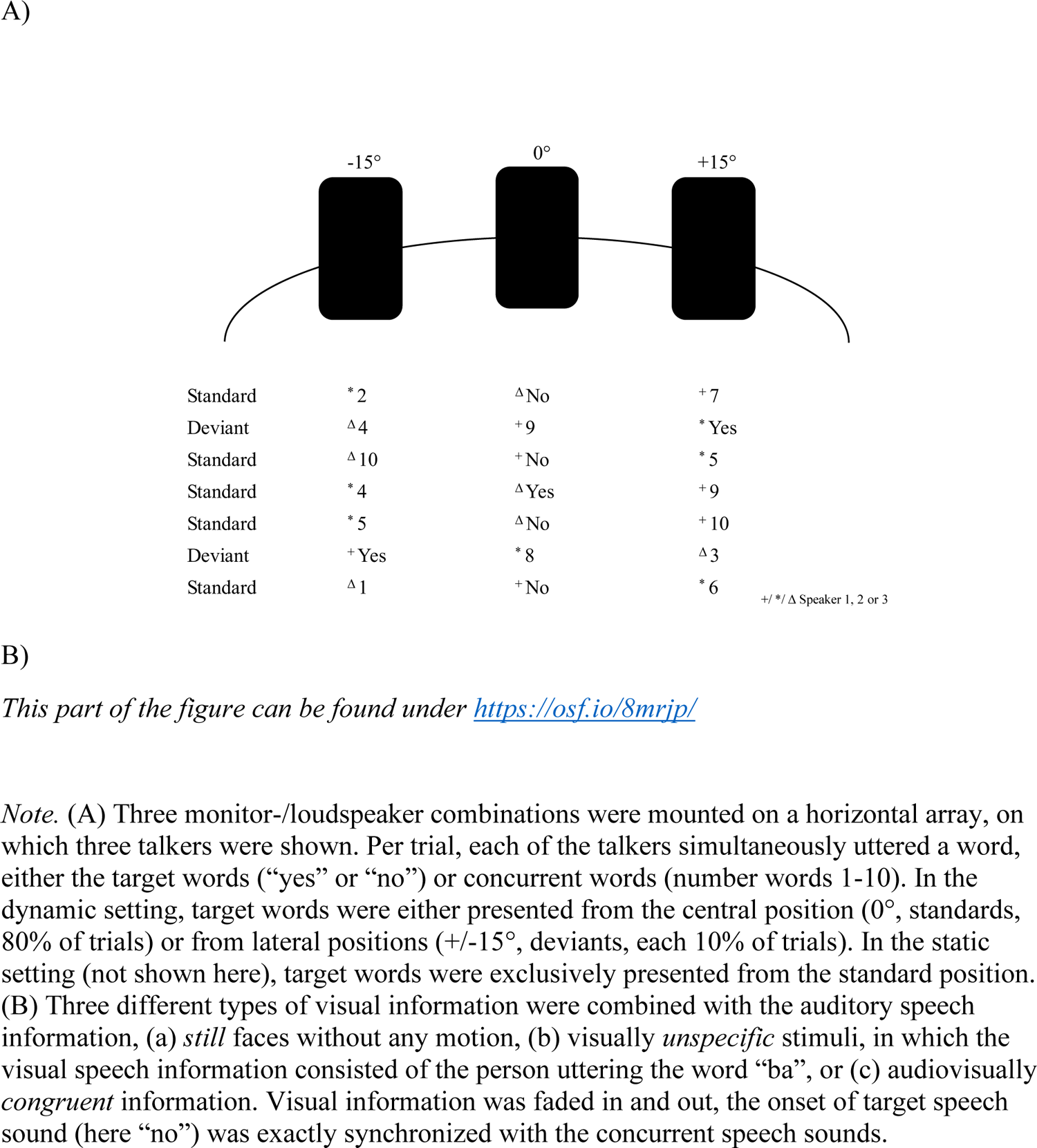
Experimental setup and audiovisual information

As stimuli, we used short video sequences showing a talker’s face and neck in front of a light-blue background (RGB values 106, 145, 161) while uttering brief words. The words were “yes” and “no” (in English) as target stimuli, and the numbers from one to ten in German language as distractor stimuli. The target words were chosen to differ clearly in visual articulation, thus providing the participants with rich visual speech information. The talkers were three young native dialect-free German females with average fundamental voice frequencies of 169 Hz (*SE* = 2.00), 213 Hz (*SE* = 7.87) and 178 Hz (*SE* = 12.22), respectively. The duration of a word was between 475 and 705 ms (*M* = 549.67, *SE* = 7.55). The talkers were instructed to pronounce the words clearly and accentuated, but to avoid unnecessary head movements.

Videos were recorded with a 1920 × 1080-pixel resolution and a 50fps frame rate under sound-attenuated conditions. While videotaping, mono audio tracks were separately recorded at 48 kHz and a 24-bit resolution using a dynamic USB-microphone (Podcaster, RØDE, Silverwater, NSW, Australia). Recording and editing was done in Audacity(R) (version 2.3.0). Audio tracks were noise reduced by 15 dB and normalized to −6 dB. The final stimulus videos started and ended with a 500-ms fade-in and a 400-ms fade-out, respectively, with freeze-frames fading from and to black. Freeze frames were taken from the beginning and ending of the visual speech video showing the talkers with their mouth closed. Between fades and speech videos, we added a variable amount of freeze frames, resulting in all stimulus videos being exactly 2900 ms long (figure 1B). Auditory speech onset was set to 1500 ms relative to the first fade-in video frame (at 0 ms) in order to ensure a synchronous onset of auditory speech in the experiment. Visual speech started and ended with the opening and closing of the talker’s mouth, resulting in visual movements that lasted between 640 and 1540 ms (*M* = 1087.43 ms, *SE* = 42.45). The visual speech onset preceded the auditory onset by 414.29ms on average (*range* = 80 – 860 ms, *SE* = 34.73).

We further edited the videos to vary the content of visual speech information provided to the participants. We generated video clips with audiovisually congruent, visually unspecific, and no visual speech information (still face). For visually unspecific information, we additionally recorded the talkers while uttering the word “ba” and manipulated the visual speech information. Thus, while the auditory information remained unaltered, the visual “ba” was used to replace the “yes” and “no” as well as the number words in the originally congruent videos. Consequently, in the visually unspecific condition, all target and distractor stimuli contained identical visual information. The resulting audiovisual videos exactly matched the timing of audiovisual congruent videos, but without containing any visual speech information that could be used for the discrimination of the target words. For the condition without visual speech information, we replaced the audiovisually congruent videos with a freeze frame of the talkers with their mouth closed, keeping the overall structure of the video unchanged (figure 1B). Videos were rendered to MP4. During the experiment, each stimulus video was followed by a fixed ISI of 1000 ms showing a black screen.

### 2.4 Experimental Procedure

Prior to the EEG experiment, every participant went through an assessment of their sensory and cognitive abilities (see 2.2). Then, participants received written and oral instructions on the task of the EEG experiment. In a two-alternative forced-choice task, the participants had to indicate as fast and accurate as possible which target word (“yes” or “no”) they heard. For this, one of the two buttons on a keypad had to be pressed with the index or middle finger of the dominant hand; the left button being assigned to “yes” (50 % of trials), the other one to “no”. Each trial consisted of the simultaneous audiovisual presentation of the three talkers, each uttering a word, always presenting one of the two target words among two distractor words (i.e., German numerals one to ten). Within a trial, distractors always differed from each other. There were no non-target trials. Trial order, target talkers and talker locations followed a pseudo-randomized order. In total, 900 trials were presented in 6 blocks. Half of those blocks were *static* blocks consisting of standard trials, in which the target words were presented exclusively from the central location. The other half were *dynamic* blocks, consisting of 80 % standard trials and 20 % deviant trials, in which the target word was either presented from the left (10 %) or right (10 %) location. A deviant trial was always followed by at least two and at most eight (on average 3.92) standard trials. Static and dynamic blocks comprised 120 and 180 trials, respectively. Audiovisually congruent, visually unspecific, and no visual speech information was presented blockwise in separate task conditions, resulting in three static and three dynamic blocks, respectively. Blockwise presentation of the audiovisual conditions was chosen with regard to the present literature (e.g. Klucharev et al., 2003; Winneke & Phillips, 2011), giving the subjects the opportunity to adjust to the information presented and thus to better elaborate the differences in audiovisual processing between the conditions. The order of static and dynamic blocks as well as audiovisual conditions was counterbalanced across participants to avoid sequence effects. Before a new block started, the participant was informed about the upcoming audiovisual condition (audiovisually congruent, visually unspecific, or no visual speech information) and whether switches of the target talker could occur or not (static or dynamic block). Between every block, participants could have a short self-paced rest, no feedback was given during the experiment. Participants were instructed to direct their gaze at the central monitor at all times and to avoid eye movements.

### 2.5 Lipreading assessment

To assess the participants’ ability to lipread, we constructed a short lip-reading assessment that followed the experimental procedure. We recorded a single female talker uttering 30 randomly assembled five-word sentences in German taken from the Oldenburger Satztest (Wagener, Kühnel, & Kollmeier, 1999). A sentence was always built the same way and consisted of a name, verb, number, adjective, and noun (e.g. “Kerstin took two old knives”). The sentences were recorded using the same equipment and setup that was used for creating the stimulus material for the EEG experiment, always starting and ending with the talker’s mouth closed. Audio tracks were removed and a 1000-ms fade with freeze frames from black was added at the beginning and end of the video using the software Shotcut.

Additional freeze frames were added prior to video onset to achieve a theoretical sound onset at 2000 ms for each video. Lip movement duration was between 2940 and 4180 ms. All videos were arranged in a presentation, presented on a MacBook positioned on the table in front of the participant. The assessment started with a detailed oral and on-screen instruction and the presentation of five example videos. After each video, two tests followed – a *cued* and a *recognition* test. For the cued test, two words were displayed on the computer screen before the video started, one of which being included in the following sentence. After watching, the participants were instructed to speak out loud which one of the two words they had seen.

Then, in the recognition test, a single word was displayed, and the participants had to state, if that word had been part of the sentence or not. For analysis, we calculated a sum score for each test based on correct answers, resulting in a possible score between 0 and 30 points. Younger participants correctly answered 23.96 (*range* = 12-30, *SE* = 0.99) of the cued questions and 17.65 (*range* = 13-27, *SE* = 0.68) of the recognition questions. Older participants correctly answered an average of 23.30 cued questions (*range* = 15-29, *SE* = 0.98) and 17.35 (*range* = 12-22, *SE* = 0.60) of the recognition questions. There were no group differences in neither the cued, *V* = 194, *p* = .519, *r* = .10, nor the recognition task, *t*(38.64) = −1.03, *p* = .310, *g* = −0.31, indicating similar lipreading performance in both age groups despite worse visual abilities in the older participants. Consequently, we did not further include the test in our analysis.

### 2.6 EEG recording and preprocessing

For continuous EEG recordings, we used a 64 Ag/AgCl electrode cap (BrainCap; Brainvision, Gilching, Germany); the electrodes were distributed across the scalp according to the extended international 10-20 system. The signal was recorded with a sampling rate of 1000 Hz (QuickAmp DC Amplifier, Brainvision, Gilching, Germany). Impedance was kept below 10 kΩ. uring the recording, the electrodes AFz and FCz were used as ground and reference electrodes, respectively.

For data preprocessing and analysis, we used MATLAB (2019a) and the open-source toolboxes EEGLAB (14-1-2b, Delorme & Makeig, 2004) and ERPLAB (v7.0, Lopez-Calderon & Luck, 2014). All data were filtered using a 0.5 Hz high-pass filter (6601-point FIR filter, 0.5 Hz transition bandwidth, 0.25 Hz cut-off frequency) and a 30 Hz low-pass filter (441-point FIR filter, 7.5 Hz transition bandwidth, cut-off frequency 33.75 Hz). To analyze the CNV-like slow wave prior to sound onset, we separately filtered the data with a 0.1 Hz high-pass filter (33001 FIR filter, 0.1 Hz transition band width, 0.05 Hz cutoff frequency) to avoid artifacts due to too strong high-pass filtering (as discussed by Tanner, Morgan-Short, & Luck, 2015). The data were re-referenced to the average of all electrodes and segmented into 2600-ms epochs covering periods from −1600 to 1000 ms relative to the auditory speech onset. The first 100 ms of the epoch served as baseline period (figure 1B). For artifact rejection, we applied independent component analysis (ICA). To reduce computation time, ICA-decomposition was based on a subset of data that contained only every second trial and was down-sampled to 250 Hz. The resulting ICA weights were then transferred back onto the ‘original’ dataset, containing all trials with a sampling rate of 1000 Hz. Artifacted components containing general discontinuities, eye blinks and eye movements where identified and subtracted from the data using the automatic artifact detection algorithm ADJUST (Mognon, Jovicich, Bruzzone, & Buiatti, 2011). Moreover, dipoles were fitted to a spherical head model using EEGLAB’s function dipfit. Components with a residual variance above 40 % in the dipole solution were subtracted from the data. Finally, we additionally removed all trials with extreme fluctuations above 1000 Hz, using the built-in EEGLAB auto-rejection function (prob. threshold: 3 std.). On average, 21.43 ICs were rejected (*range* = 8 - 38) and 15.54 % of trials were excluded per subject (*range* = 7.44 - 26.33 %). For further analyses, only trials with correct responses were included, which led to an additional exclusion of an average of 40.30 trials per subject (being 0.56 to 21.22 % of the original amount of trials). After preprocessing, the data were down-sampled to 500 Hz for further analyses.

### 2.7 Data Analysis

We performed statistical analysis using R (v3.6.3) in RStudio (RStudio Team, 2020, version 1.2.5042). For the performed analyses of variance (ANOVAs) and subsequent post-hoc pairwise *t*-tests, partial eta squared *η_p_²* (Cohen, 1973) and Hedges’s *g,* (i.e., corrected Cohen’s *d* to account for bias due to smaller sample sizes, Fritz, Morris, & Richler, 2012) were calculated as effect size measures, respectively. Multiple comparisons were corrected using FDR (Benjamini & Hochberg, 1995). Those parametric tests were conducted, if the Shapiro-Wilk test yielded *p* > .05 and thus indicated approximately normally distributed data (Field, Miles, & Field, 2012). In case of a violation of the normality assumption, non-parametric analyses were chosen, that is, rank-based ANOVA-type statistics (as provided by the R-package nparLD; Noguchi, Gel, Brunner, & Konietschke, 2012). Note, that except for the whole-plots factor age group, only the nominator degrees of freedom will be reported in case of ANOVA-type statistics, since the denominator is set to infinite. Also note, that *F-* values are not necessarily comparable to parametric *F*-values (Noguchi et al., 2012). For non-parametric post-hoc calculations, we used the Wilcoxon signed-rank test (Wilcoxon, 1945) with *r* as an effect size measure (Fritz et al., 2012).

#### 2.7.1 Behavioral analyses

As behavioral measures, we analyzed accuracy and response times. Response times were defined as the time interval between the end of video fade-in and a button press. We chose this early timepoint to avoid negative response times, which could occur in principle, since a response before sound-onset was possible due to the preceding visual information. It should be noted, however, that in the way the stimulus videos were constructed, the end of the fade-in always preceded the sound-onset by 1000ms. Only trials with correct response were taken into the analysis, response times below three standard deviations from the mean were excluded as pre-mature responses. Separate analyses were performed for the *static* and *dynamic* settings: For the static setting, in which only standard trials (i.e., with targets presented from the central position) were presented, a 2 (young vs. old) × 3 (congruent vs. unspecific vs. still face visual information) repeated-measures ANOVA was conducted. For the dynamic setting, with both standard and deviant trials (i.e., with targets presented from central and lateral positions), we performed a 2 (young vs. old) × 3 (congruent vs. unspecific vs. still face visual information) × 2 (standard vs. deviant target position) repeated-measures ANOVA.

#### 2.7.2 ERP analyses

2.7.3 For ERP analyses, we computed grand-averages across participants, separately for the two age groups, the three audiovisual conditions, and the static and dynamic blocks. In addition, for dynamic blocks, separate averages were computed for deviant and standard trials. However, to avoid post-deviant modulations of the waveforms (e.g., Getzmann & Wascher, 2016), standard trials immediately following a deviant (i.e., the two trials post-deviant presentation) were not considered in the analysis of dynamic blocks. All data were collapsed across both target stimuli and target talkers. ERP data, that is, correct trials, were averaged across a cluster of frontocentral electrodes that included the electrodes Cz, FCz, Fz, FC1 and FC2. To analyze the amplitudes of P1, N1, P2 and N2, we determined the maximum or minimum peak in the grand-average waveforms in a previously defined time window (P1: 20 – 120 ms, N1: 75 – 175 ms, P2: 150 – 250 ms, N2: 250 – 350 ms) at the above-mentioned electrodes. Around the detected peaks, 20-ms time windows were used to compute the mean amplitude. In order to account for the broad negative, CNV-like deflection, which peaked immediately before sound onset and which appeared to differ between audiovisual conditions (cf., figures 3 and 6), we analyzed mean-amplitude-to-mean-amplitude measures of the P1, N1, P2 and N2 components. We calculated the absolute difference between the mean amplitudes of two respective components. For the analysis of the CNV, we calculated the mean amplitudes at the same cluster of frontocentral electrodes within the time window 100 ms prior to the auditory speech onset. Data for the static setting were analyzed in a 2 (young vs. old) × 2 (congruent vs. unspecific visual information) repeated-measures ANOVA. The still face condition was not included in the ERP analyses, because ERP differences between stimuli with still face in comparison to audiovisually congruent and visually unspecific information could be confounded by sensory differences of the three conditions. In particular, ERP differences between the audiovisual congruent and the still face condition could be found due to physical stimulus differences (moving faces vs. still faces) and less to the content-specific difference, which should be apparent when comparing the audiovisual congruent and the unspecific conditions (both including moving faces). For the dynamic setting, again, we calculated repeated-measures ANOVAs with the additional factor target position (standard vs. deviant target position).

#### 2.7.4 Supplementary analyses

We conducted several additional analyses (see Supplementary Material). First, in order to assess the extent to which behavioral performance depended on basic unisensory and cognitive abilities, we calculated pearman’s correlations (Spearman & Spearman, 1904) between the unisensory and cognitive abilities (i.e., visual acuity, hearing level, MoCA score, and working memory capacity) and the behavioral measures (i.e., accuracy and response time). Second, to investigate whether standard trials were processed and reacted to differently depending on the setting (i.e., static or dynamic), we conducted a 2 (young vs. old) × 3 (congruent vs. unspecific vs. still face visual information) × 2 (static vs. dynamic setting) repeated-measures ANOVA for accuracy and response times. For the ERP measures P1-N1, N1-P2, P2-N2, we conducted a similar analysis, leaving out the still face condition. Finally, to investigate ERP latency differences between the experimental conditions, we analyzed latency effects for P1, N1, P2, and N2 in a 2 (young vs. old) × 2 (congruent vs. unspecific visual information) repeated-measures ANOVA for the static setting, and in a 2 (young vs. old) × 2 (congruent vs. unspecific visual information) × 2 (standard vs. deviant target position) repeated-measures ANOVA for the dynamic setting.

## 3. Results

The analysis of the behavioral and ERP data is divided into two parts: we first looked at the effects of age and visual information in the *static* setting, and then in the *dynamic* setting, where we compared standard and deviant trials.

### 3.1 Static setting

#### 3.1.1 Behavioral data Accuracy

In figure 2A, behavioral data are visualized, showing accuracy and response times depending on age group and visual information. Older participants reacted less accurate on the task (*M_old_* = 94.60 %, *SE_old_* = 0.93; *M_young_* = 97.39 %, *SE_young_* = 0.39; figure 2A; table 1). The rate of correct responses also depended on the visual information, an effect which was modulated by an interaction with age (cf. figure 2A). Post-hoc comparisons showed that older participants performed more accurately with audiovisually congruent stimuli compared to visually unspecific information, *V* = 176, *p_adj_* = .006, *r* = .74, and still face stimuli, *V* = 111, *p_adj_* = .012, *r* = .70; performance in the last two conditions did not differ, *V* = 110, *p_adj_* = .576, *r* = .25. However in younger participants, no differences in accuracy between the content of visual information was shown (congruent vs. still *t*(21) = 0.72, *p_adj_* = .576, *g* = 0.15; congruent vs. unspecific, *V* = 66, *p_adj_* = .754, *r* = .0176; still vs unspecific, *t*(21) = −0.70, *p_adj_* = .576, *g* = - 0.14).

**Figure 2.**
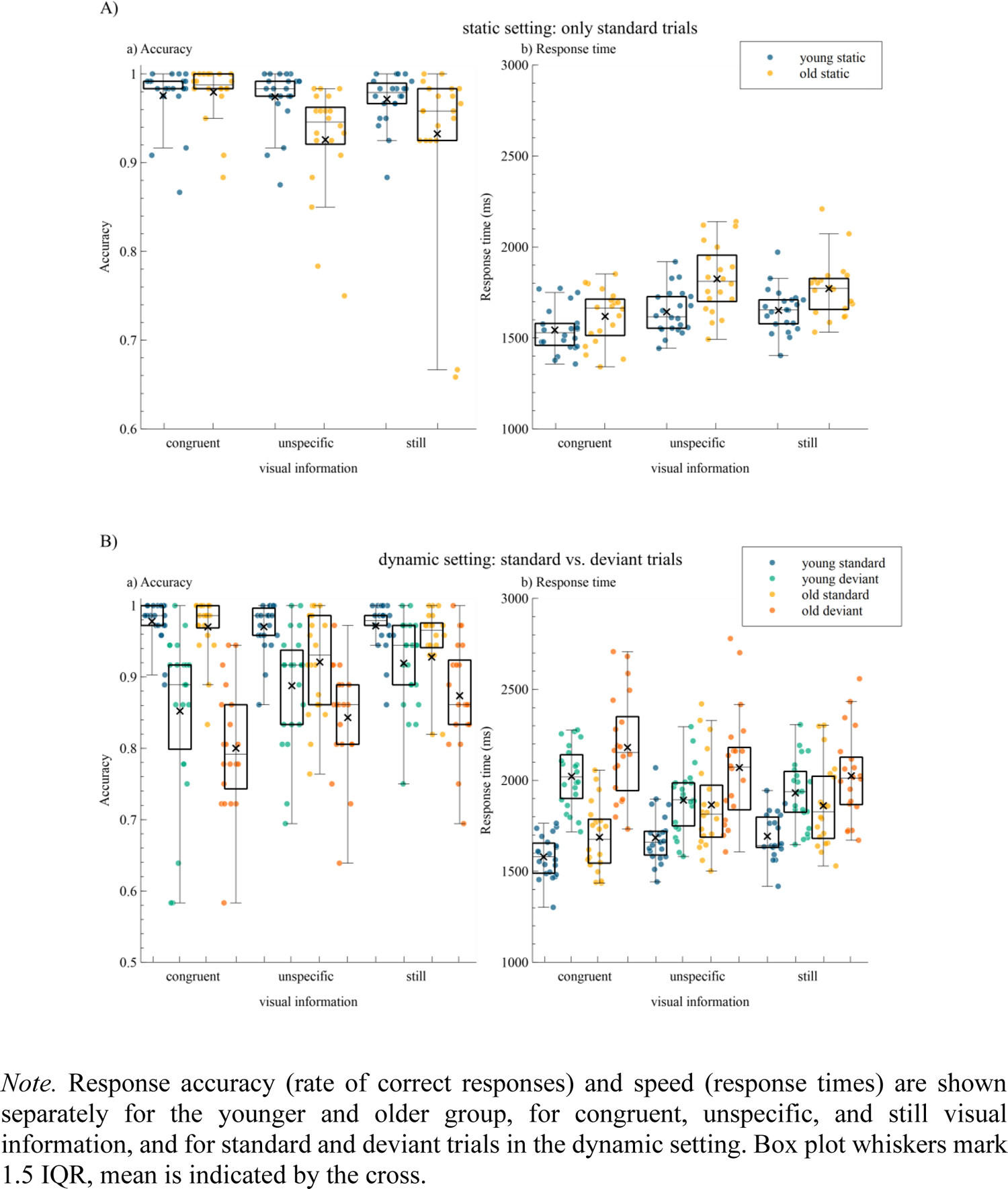
Behavioral results for the (A) static and (B) dynamic setting

**Table 1.** ANOVA results for trials in static setting

| conditions | Accuracy<sup>1</sup> | Response time<sup>2</sup> |  | P1-N1<sup>1</sup> | N1-P2<sup>3</sup> |  | P2-N2<sup>3</sup> |  | CNV<sup>3</sup> |  |
| --- | --- | --- | --- | --- | --- | --- | --- | --- | --- | --- |
| | *F* | *F* | $\eta_p^2$ | *F* | *F* | $\eta_p^2$ | *F* | $\eta_p^2$ | *F* | $\eta_p^2$ |
| age | **4.83\*** | **9.32\*\*** | .19 | **4.70\*** | 3.61<sup>T</sup> | .08 | **13.76\*\*** | .26 | 0.20 | <.01 |
| visual information | **14.26\*\*\*** | **54.10\*\*\*** | .57 | **7.80\*\*** | 1.36 | .03 | **33.73\*\*\*** | .46 | **9.95\*\*** | .20 |
| age × visual information | **9.39\*\*\*** | **5.69\*\*** | .12 | **5.05\*** | 0.01 | <.01 | 1.52 | .04 | 0.02 | <.01 |

*Notes.* 2 (young vs. old) × 2 (congruent vs. unspecific visual information) repeated-measures ANOVA in the static setting. For accuracy and response time, the additional condition *still faces* was included for the factor of visual information.
<sup>1</sup> ANOVA-type statistic; except for whole-plots factors, denominator degrees of freedom are infinite, thus reporting only nominator degrees of freedom.
Degrees of freedom: visual information, age × visual information = 1.95 (accuracy), 1.00 (P1-N1); age = 1,39.94 (accuracy), 1,37.40 (P1-N1).
<sup>2</sup> Degrees of freedom: Age = 1,40; visual information, age × visual information = 2, 80.
<sup>3</sup> Degrees of freedom: 1,40 for all factors.
<sup>T</sup> $p < .10$ , \* $p < .05$ , \*\* $p < .01$ , \*\*\* $p < .001$ .

#### Response times

In the older group, higher response times were found compared to the younger group (*M_old_* = 1738.22 ms, *SE_old_* = 24.06, *M_young_* = 1612.83 ms, *SE_young_* = 16.01; figure 2A; table 1). Additionally, the analysis revealed a main effect for visual information, which interacted with the age group. However, post-hoc tests indicated that for both the younger and older group, response times following audiovisually congruent information were lowest compared to visually unspecific information, old: *t*(19) = −6.80, *p_adj_* < .001, *g* = −1.46, young: *t*(21) = −5.70, *p_adj_* < .001, *g* = −1.17, and still faces, old: *V* = 0, *p_adj_* < .001, *r* = .88, young: *V* = 16, *p_adj_* < .001, *r* = .77. In both groups, there was no difference between visually unspecific and still face stimuli, old: *t*(19) = −1.75, *p_adj_* = .115, *g* = −0.36, young: *t*(21) = 0.85, *p_adj_* = .403, *g* = 0.18.

#### 3.1.2 ERP data

Looking at the grand-averaged waveforms at frontocentral scalp locations, the onset of the (still) faces of the talkers (at −1500 ms, figure 3) elicited a sequence of deflections which was related to genuine visual processing and not analyzed here. The onset of the auditory information (at 0 ms, i.e., the time-locking event) elicited a prominent pattern of deflections reflecting the P1, N1, P2, and N2 components, which was preceded by a broad, CNV-like negativity. This series of deflections will be further analyzed and described below. Note that the CNV analysis was conducted on 0.1 Hz data (for the ERP waveform on 0.1 Hz, see Supplementary Material, figures S1).

**Figure 3.**
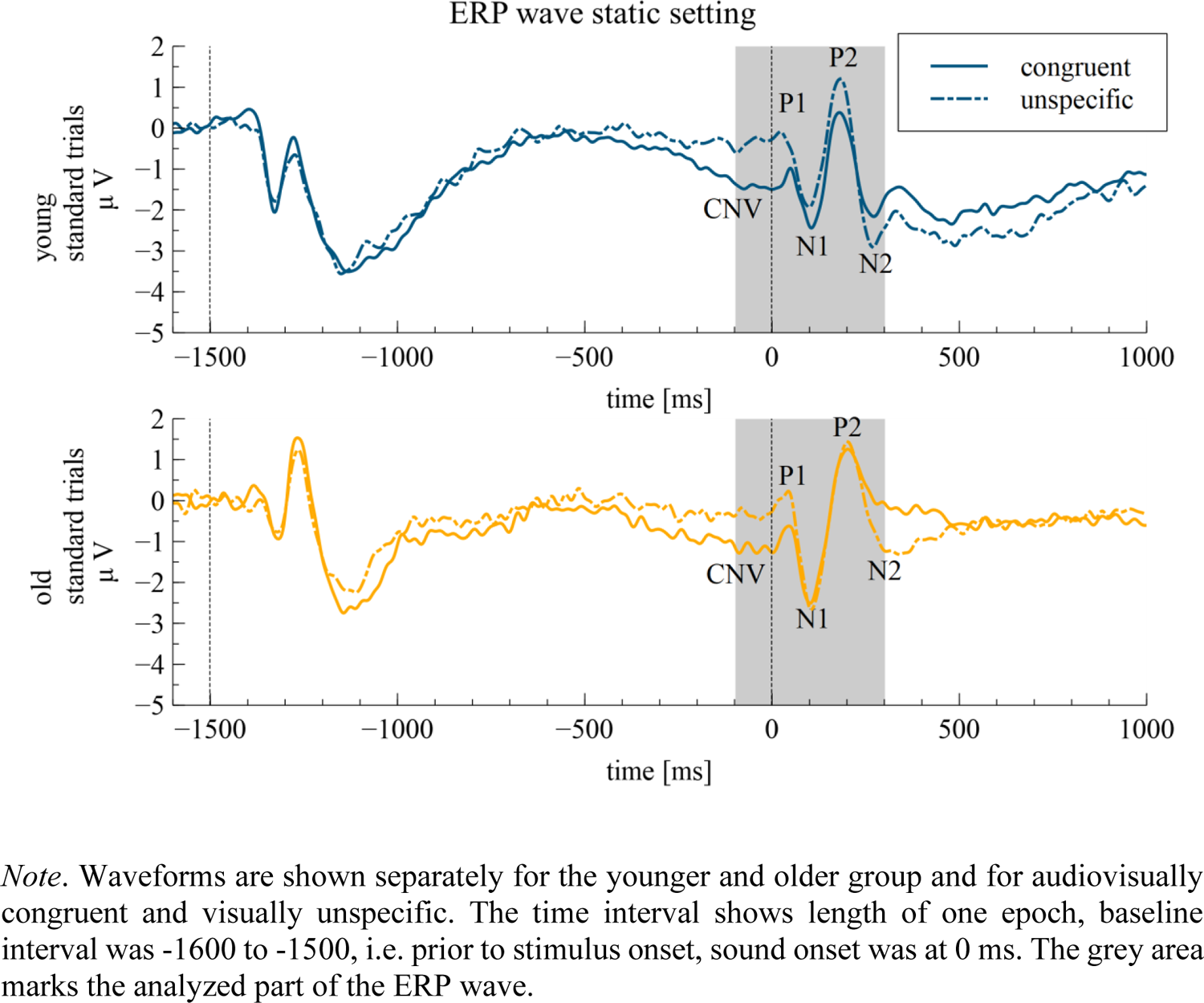
ERP waveforms at electrode cluster Cz, FCz, Fz, FC1 and FC2 in static setting

#### P1-N1

In the early processing stage, reflected by the P1 to N1 amplitude measure, the analysis revealed larger amplitude measures for the older (*M* = 2.31 µV, *SE* = 0.24) compared to the younger group (*M* = 1.52 µV, *SE* = 0.15, figure 4, table 1). Moreover, amplitude measures were smaller for audiovisually congruent (*M* = 1.67 µV, *SE* = 0.18) compared to visually unspecific stimuli (*M* = 2.12 µV, *SE* = 0.21). An interaction with the age group revealed that the difference between the visual conditions was only present in older, *V* = 26, *p_adj_* = .004, *r* = .66, but not in younger adults, *t*(21) = −0.13, *p_adj_* = .897, *g* = −0.03.

**Figure 4.**
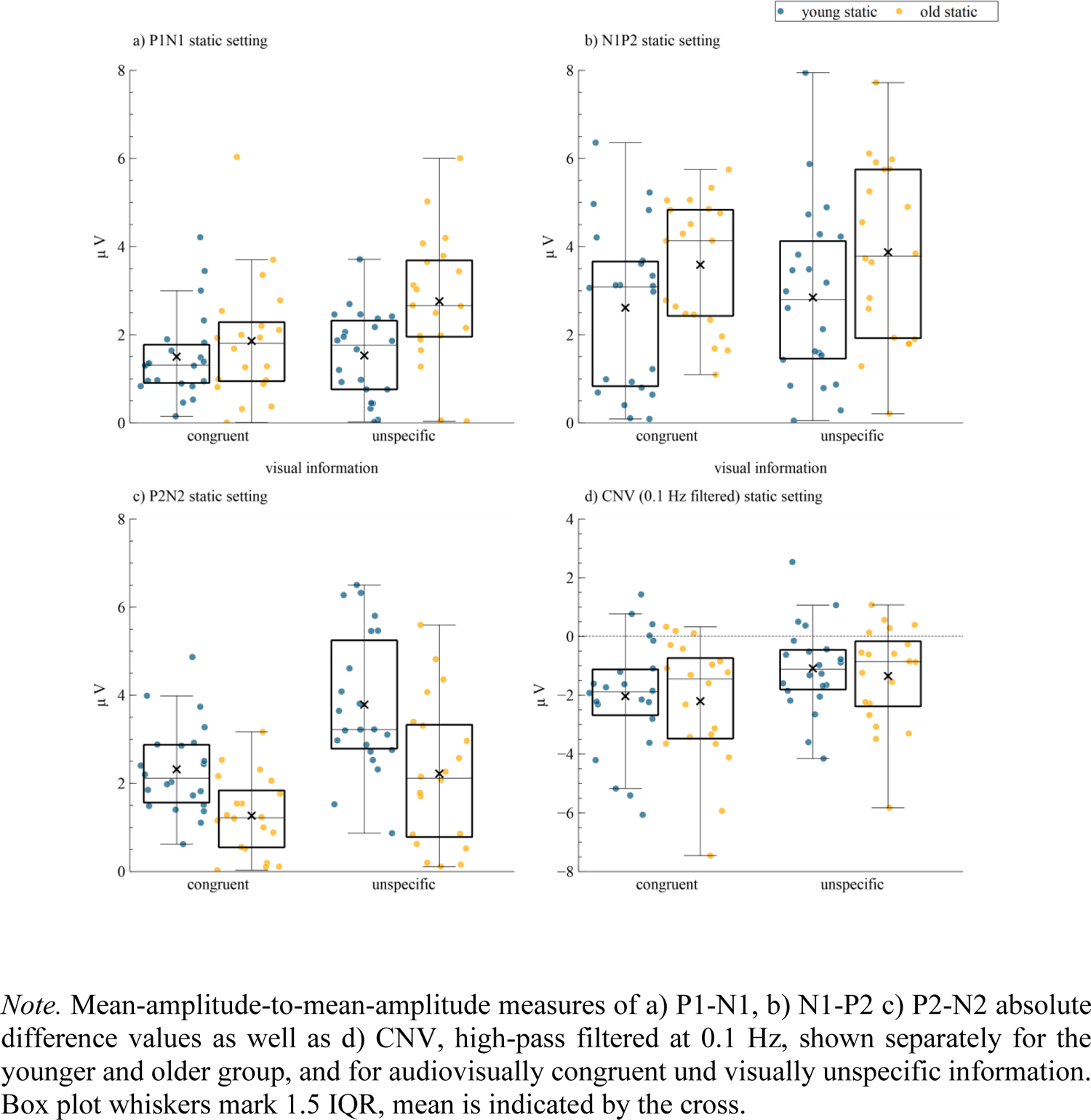
ERP amplitudes in static setting

#### N1-P2

For the N1 to P2 amplitude measure, analysis did not reveal any significant effects (figure 4, table 1).

#### P2-N2

For the later processing stage, as reflected by P2 to N2 amplitude measures, we found smaller amplitude measures in the older (*M* = 1.74 µV, *SE* = 0.22) compared to the younger group (*M* = 3.05 µV, *SE* = 0.23, figure 4, table 1). Amplitude measures for audiovisually congruent stimuli (*M* = 1.82 µV, *SE* = 0.17) were significantly smaller than for visually unspecific stimuli (*M* = 3.04 µV, *SE* = 0.28).

#### CNV

The CNV amplitude was more pronounced for audiovisually congruent stimuli (*M* = −2.11 µV, *SE* = 0.31) than for visually unspecific stimuli (*M* = −1.21 µV, *SE* = .24, figure 4 and 5, table 1), while no difference between age groups or interaction occurred.

**Figure 5.**
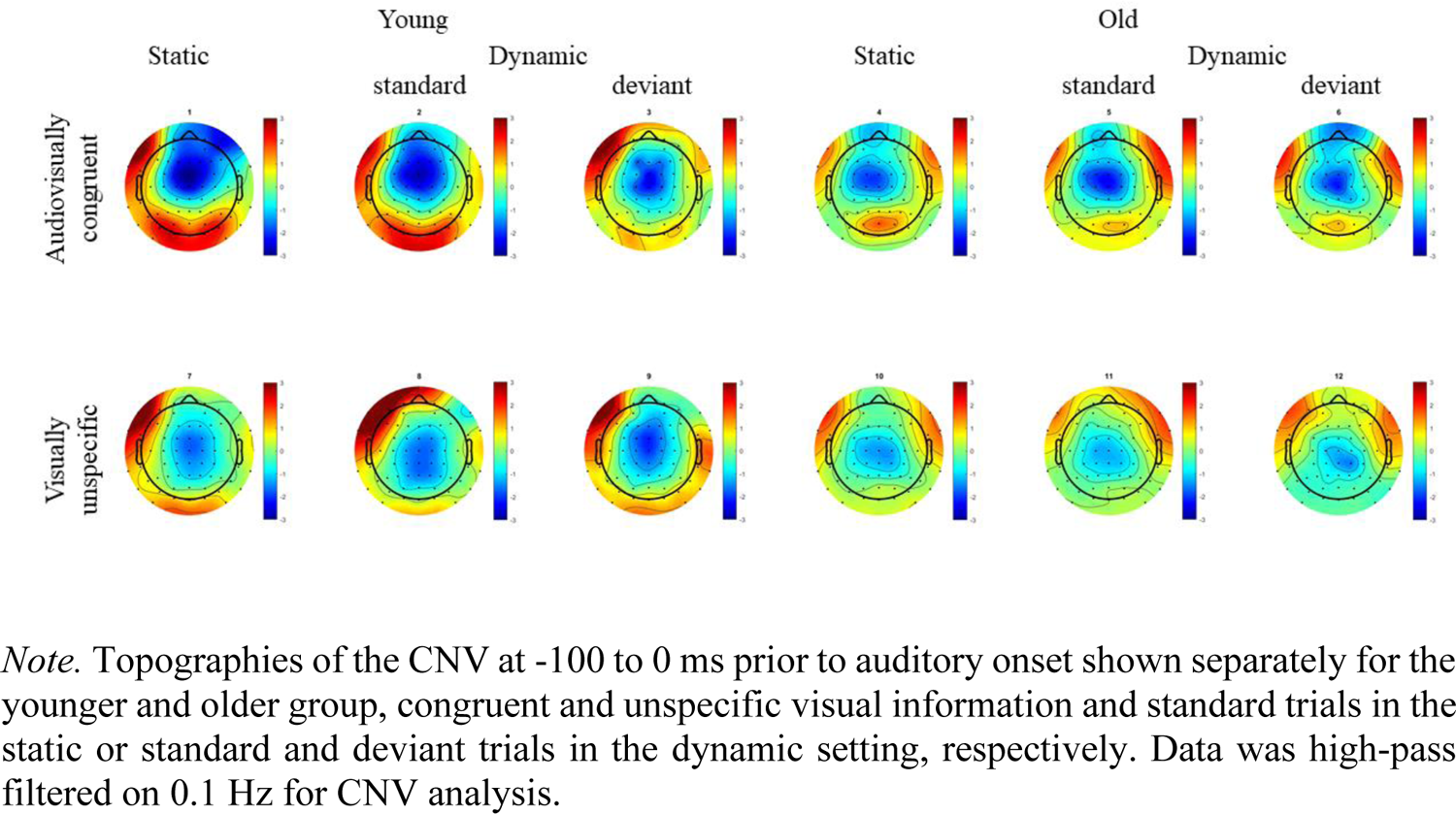
Topographies of the CNV at 100 to 0 ms

### Summary: static setting

Overall, the behavioral data of the static setting indicated that younger adults showed a better performance on the task than older adults. Additionally, performance was better with audiovisually congruent information relative to unspecific information and still faces, especially in the older group, which suggests a facilitation of the task when the seen lip movement was congruent to the auditory speech input. ERP amplitude measures were enhanced in the older group in early processing (P1-N1), but decreased in late processing (P2-N2). For both early and late processing, audiovisually congruent information resulted in decreased amplitudes compared to visually unspecific information. Additionally, we found a more pronounced CNV amplitudes for audiovisually congruent stimuli.

### 3.2 Dynamic setting: Standard vs. deviant stimuli

#### 3.2.1 Behavioral data

##### Accuracy

Within the dynamic setting, the rate of correct responses was lower in older, than younger participants (*M_old_* = 88.92 %, *SE_old_* = 0.90; *M_young_* = 92.98 %, *SE_young_* = 0.72; figure 2B; table 2). Accuracy was also higher in standard, compared to deviant trials (*M_std_* = 95.72%, *SE_std_* = 0.58; *M_dev_* = 86.38 %, *SE_dev_* = 0.82). In addition, we found an interaction of visual information and target location. Post-hoc comparisons revealed that within standard trials, accuracy was higher for audiovisually congruent (*M* = 97.42 %, *SE* = 0.55) than for visually unspecific stimuli (*M* = 94.68 %, *SE* = 0.93; *V* = 438.5, *p_adj_* = .010, *r* = .40) and still faces (*M* = 95.07 %, *SE* = 1.34; *V* = 352.5, *p_adj_* = .009, *r* = .43), with no differences between last two, *V* = 334, *p_adj_* = .994, *r* = .002. In contrast, within deviant trials, accuracy was lower in audiovisually congruent (*M* = 82.73 %, *SE* = 1.66) compared to visually unspecific stimuli (*M* = 86.64 %, *SE* = 1.26; *t*(41) = −2.51, *p_adj_* = .024*, g* = −0.38) and still faces (*M* = 89.75 %, *SE* = 1.13;*t*(41) = −5.78, *p_adj_* =< .001, *g* = −0.88). Also, accuracy for visually unspecific stimuli was significantly lower than for still faces, *V* = 344.5, *p_adj_* = .028, *r* = .33.

**Table 2.** ANOVA results for standard and deviant trials in the dynamic setting

| conditions | Accuracy<sup>1</sup> | Response time<sup>2</sup> |  | P1-N1<sup>3</sup> |  | N1-P2<sup>1</sup> | P2-N2<sup>1</sup> | CNV<sup>1</sup> |
| --- | --- | --- | --- | --- | --- | --- | --- | --- |
| | *F* | *F* | $\eta_p^2$ | *F* | $\eta_p^2$ | *F* | *F* | *F* |
| age | **8.78\*\*** | **6.62\*** | .14 | 4.07<sup>T</sup> | .09 | 3.32 | **5.89\*** | 0.36 |
| visual information | 2.40<sup>T</sup> | 0.25 | .01 | **17.64\*\*\*** | .31 | **9.71\*\*** | **33.91\*\*\*** | **16.49\*\*\*** |
| target location | **173.54\*\*\*** | **199.72\*\*\*** | .83 | **4.51\*** | .10 | 0.11 | **17.44\*\*\*** | 2.40 |
| age × visual information | 0.80 | 1.40 | .03 | **6.85\*** | .15 | 0.99 | 0.79 | 0.04 |
| target location × visual information | **17.15\*\*\*** | **111.40\*\*\*** | .74 | 0.40 | .01 | 1.81 | 1.33 | **4.14\*** |
| age × target location | 0.21 | 0.04 | < .01 | 1.54 | .04 | 1.80 | 0.03 | **4.48\*** |
| age × visual information<br>× target location | 1.96 | **4.84\*** | .11 | 3.83<sup>T</sup> | .09 | 0.06 | 0.81 | 0.03 |

Notes. 2 (young vs. old) × 2 (congruent vs. unspecific visual information) × 2 (standard vs. deviant target location) repeated-measures ANOVA in the dynamic setting. For accuracy and response time, the additional condition *still faces* was included for the factor of visual information.
<sup>1</sup> ANOVA-type statistic, except for whole-plots factors, denominator degrees of freedom are infinite, thus reporting only nominator degrees of freedom.
Degrees of freedom: age = 1, 39.23 (accuracy), 1.00 (N1-P2, P2-N2, CNV); visual information, age × visual information = 1.74 (accuracy), 1.00 (N1-P2, P2-N2, CNV); target location, group × target location = 1.00; target location × visual information, age × target location × visual information = 1.96 (accuracy), 1.00 (N1-P2, P2-N2).
<sup>2</sup> Degrees of freedom: Age, target location, age × target location = 1, 40, visual information, age × visual information, visual information × target location, age × visual information × target location = 2, 80.
<sup>3</sup> Degrees of freedom: *df* = 1,40 for all factors.
<sup>T</sup> $p < .10$ , \* $p < .05$ , \*\* $p < .01$ , \*\*\* $p < .001$ .

##### Response Times

Response times were overall larger in older than younger participants (*M_old_* = 1948.25ms, *SE_old_* = 27.40; *M_young_* = 1800.26 ms, *SE_young_* = 19.12), and in deviant than standard trials (*M_dev_* = 2016.63 ms, *SE_dev_* = 22.14; *M_std_* = 1724.83 ms, *SE_std_* = 18.40; figure 2B, table 2). There was also an interaction of visual information and target location. In standard trials, response times in audiovisually congruent stimuli were lower (*M* = 1630.71 ms, *SE* = 25.21) compared to visually unspecific (*M* = 1770.91 ms, *SE* = 34.61, V = 11, *p_adj_* < .001, *r* = .85) and still face stimuli (*M* = 1772.87 ms, *SE* = 30.67, *t*(41) = −9.86, *p_adj_* < .001, *g* = −1.49), with no difference between the two latter, *V* = 535, *p_adj_* = .363, *r* = .16. In contrast, response times in deviant trials were higher with audiovisually congruent (*M* = 2097.45 ms, *SE* = 37.73), compared to both visually unspecific (*M* = 1976.68 ms, *SE* = 41.15, *t*(41) = 5.67, *p_adj_* < .001, *g* = 0.86) and still face stimuli (*M* = 1975.75ms, *SE* = 33.59, *t*(41) = 5.74, *p_adj_* < .001, *g* = 0.87). Again, the two latter did not differ, *V* = 494, *p_adj_* = .603, *r* = .08.

Finally, there was a three-way interaction of age, visual information, and target location. Separate analyses for both age groups showed a similar pattern, with a significant interaction of visual information and target location, old: *F*(38,2) = 56.03, *p_adj_* < .001, *η_p_²* = .75, young: *F*(42,2) = 59.48, *p_adj_* < .001, *η_p_²* = .74. Post-hoc comparisons in the older adults showed smaller response times in standard trials, when comparing audiovisually congruent with visually unspecific information, *t*(19) = −4.99, *p_adj_* < .001, *g* = −1.07, and still faces, *t*(19) = −7.53, *p_adj_* < .001, *g* = −1.62. In deviant trials, larger response times followed audiovisually congruent compared to visually unspecific, *t*(19) = 3.61, *p_adj_* = .003, *g* = 0.78, and still face stimuli, *t*(19) = 4.91, *p_adj_* < .001, *g* = 1.05. Still face stimuli did not differ from visually unspecific stimuli in terms of response times in neither standard, *t*(19) = −0.13, *p_adj_* = .900, *g* = −0.03, nor deviant trials, *t*(19) = −1.22, *p_adj_* = .277, *g* = −0.26. In the younger adults, a similar pattern could be observed with response times being smaller in audiovisually congruent compared to visually unspecific, *t*(21) = −5.54, *p_adj_* < .001, *g* = −1.14, and still face stimuli, *t*(21) = −7.12, *p_adj_* < .001, *g* = −1.46, when shown in standard trials. The last two conditions did not differ, *t*(21) = 0.54, *p_adj_* = .641, *g* = 0.11. When deviant trials were presented, response times were larger in audiovisually congruent compared to visually unspecific, *t*(21) = 4.31, *p_adj_* < .001, *g* = 0.89, and still face stimuli, *t*(21) = 3.31, *p_adj_* = .004, *g* = 0.68. Here, also smaller response times were found in visually unspecific stimuli in comparison to still faces, *t*(21) = −2.32, *p_adj_* = .038, *g* = −0.48.

#### 3.2.2 ERP data

Looking at the grand-averaged waveforms at frontocentral scalp locations (figure 6), we can observe a similar pattern of activation as found for the static setting (figure 3). Again, prior to the onset of auditory information (0 ms) and thus prior to our components of interest, P1, N1, P2 and N2, we observed a negative, CNV-like deflection, starting about 300 ms, and peaking at approximately 100 ms prior to sound onset (Again, for the ERP waveform on 0.1 Hz, see Supplementary Material, figure S2).

**Figure 6.**
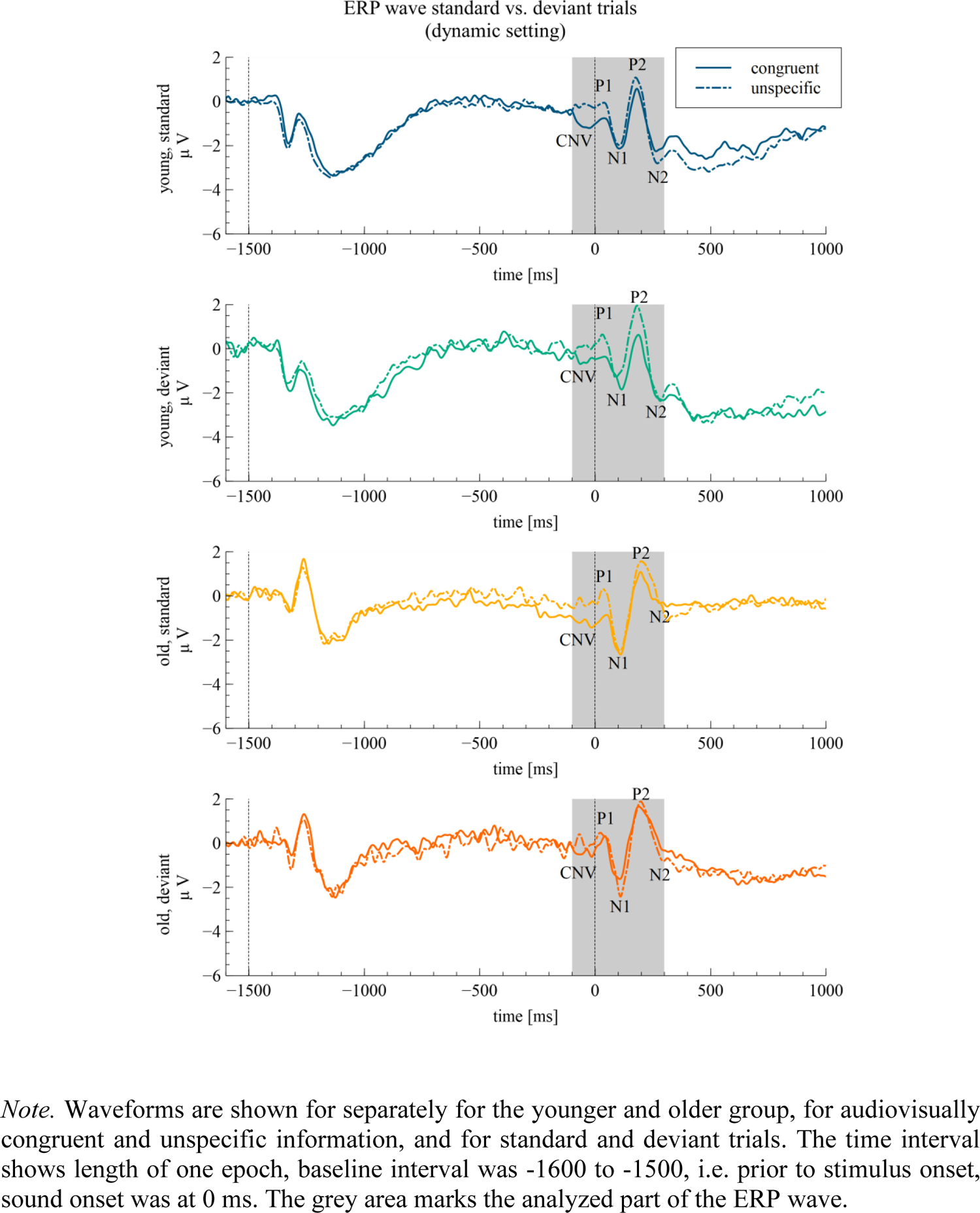
ERP waveforms at electrode cluster Cz, FCz, Fz, FC1 and FC2 in dynamic setting

**Figure 7.**
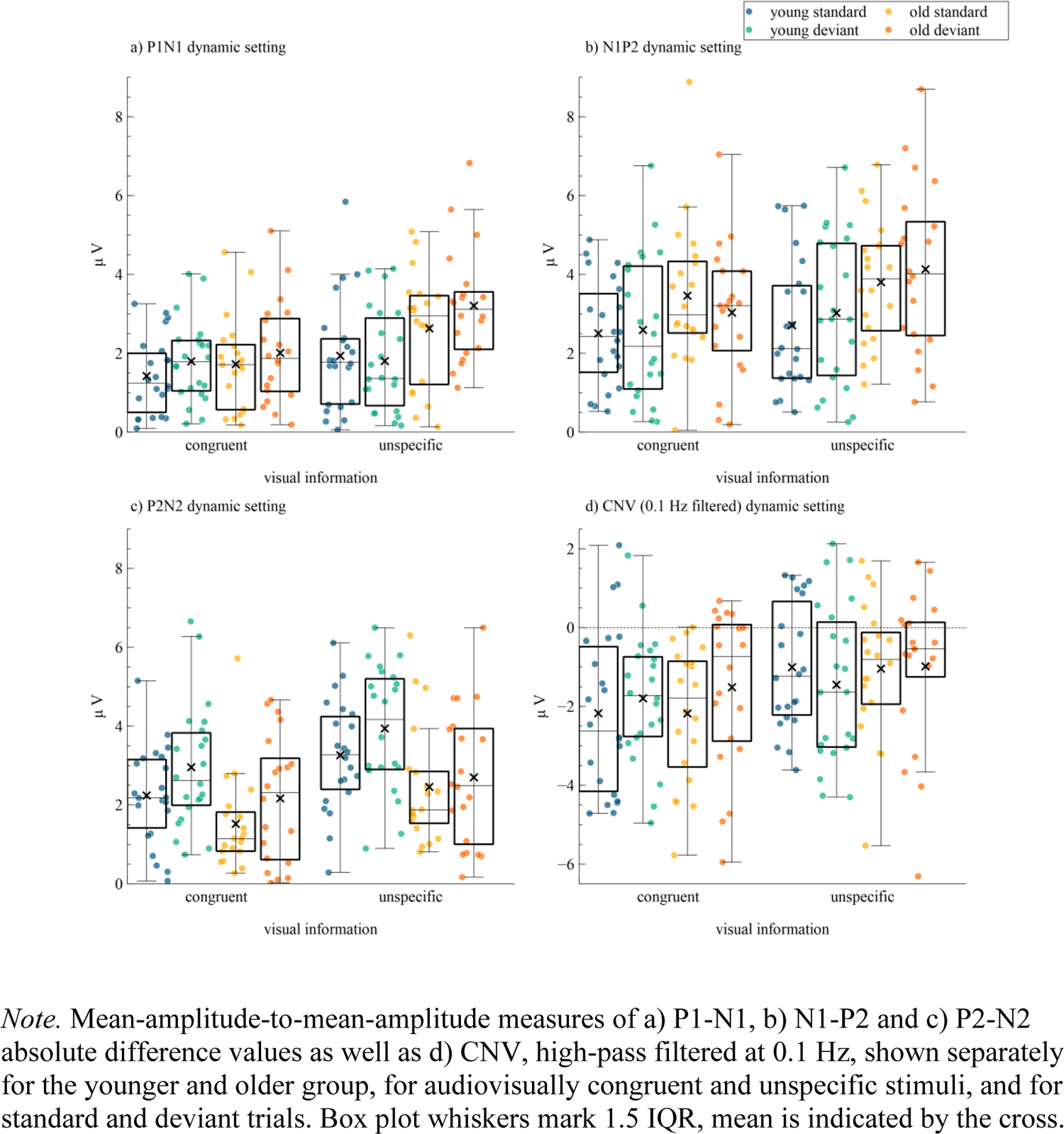
ERP amplitudes in dynamic setting

##### P1-N1

For the early processing stage, as reflected by the P1 to N1 amplitude measure, the analysis revealed that in deviant trials (*M* = 2.18 µV, *SE* = 0.15), amplitude measures were enhanced compared to standard trials (*M* = 1.92 µV, *SE* = 0.14, figure 6, table 2). Smaller amplitude measures for audiovisually congruent (*M* = 1.73 µV, *SE* = 0.12) compared to visually unspecific stimuli (*M* = 2.37 µV, *SE* = 0.16) were found. This main effect was modulated by the age group, showing that the difference between visual conditions was only present in the older, *M_congr_* = 1.86 µV, *SE_congr_* = 0.19, *M_unsp_* = 2.92 µV, *SE_unsp_* = 0.23, *t*(39) = − 6.29, *p_adj_* < .001, *g* = −0.98, but not in the younger group, *M_congr_* = 1.61 µV, *SE_congr_* = 0.15, *M_unsp_* = 1.87 µV, *SE_unsp_* = 0.20, *t*(43) = −1.33, *p_adj_* = .191, *g* = −0.20.

##### N1-P2

The analysis for the N1 to P2 amplitude measure revealed smaller amplitude measures for audiovisually congruent (*M* = 2.88 µV, *SE* = 0.19) compared to visually unspecific stimuli (*M* = 3.39 µV, *SE* = 0.21, figure 6, table 2).

##### P2-N2

For the late processing stage, as reflected by the P2 to N2 amplitude measure, we found smaller amplitude measures in the older (*M* = 2.21 µV, *SE* = 0.18) compared to the younger group (*M* = 3.10 µV, *SE* = 0.17). In deviant trials (*M* = 2.96 µV, *SE* = 0.19), amplitude measures were enhanced in comparison to standard trials (*M* = 2.39 µV, *SE* = 0.16). Additionally, amplitude measures were smaller for audiovisually congruent (*M* = 2.24 µV, *SE* = 0.16) compared to visually unspecific stimuli (*M* = 3.11 µV, *SE* = 0.18, figure 6, table 2).

##### CNV

The CNV amplitude for more pronounced for audiovisually congruent (*M* = −1.92 µV, *SE* = 0.20) compared to visually unspecific stimuli (*M* = −1.13 µV, *SE* = 0.20). This main effect was modulated by the target position, showing that this difference between the visual conditions was significant for standard trials, *t*(41) = −4.82, *p_adj_* < .001, *g* = −0.73, but not for deviant trials, *t*(41) = −1.69, *p_adj_* = .099, *g* = −0.26. Moreover, target position interacted with the age group. However, subsequent pairwise comparisons did not revealed differences between the visual conditions within the younger, *t*(43) = −0.15, *p_adj_* = .882, *g* = −0.02, and older group, *t*(39) = 1.87, *p_adj_* = .137, *g* = 0.29.

#### 3.2.3 Summary: dynamic setting

Accuracy declined and response times increased when the target stimulus was presented from a deviant position, and in both age groups these effects were stronger with audiovisually congruent stimuli than visually unspecific stimuli and still faces.

In both early (P1-N1) and late (P2-N2) processing steps, amplitude measures were enhanced in deviant compared to standard trials. Moreover, smaller amplitudes were elicited with audiovisually congruent information compared to visually unspecific information in all three measures. For the P1-N1 amplitude measure, this was only the case for the older group. Finally, we observed an amplitude reduction in the older group for the P2-N2 amplitude measures. For the CNV, larger amplitudes were found for standard trials with audiovisually congruent compared to visually unspecific information. Overall, the results in the dynamic setting are in line with those in the static setting.

## 4. Discussion

The present study investigated the processing of simultaneously presented audiovisual speech in a multi-talker scenario, comparing the neuro-cognitive mechanisms of audio-visual speech integration in younger and older adults. We varied the informational content of the visual input by presenting audiovisually congruent, visually unspecific, and still face speech stimuli. Moreover, we tested whether expected benefits of audiovisually congruent speech information (relative to incongruent and still face, that is auditory-only information) also occur under highly dynamic listening conditions. We therefore used either a static setting in which the talker of the target information was kept constant, or a dynamic setting in which the target talker occasionally switched to a deviant (lateral) position.

In both age groups, audiovisually congruent information proved to be beneficial for a faster and more accurate performance, relative to audio-only information (i.e., the still face condition), while visually unspecific information was also not particularly helpful. Overall, performance was declined in older adults, with lower accuracy rates and higher response times. Moreover, the static setting was superior to the dynamic one, in which the audiovisual benefit vanished when the target information was not presented at the expected standard location.

These behavioral results are reflected in the examined ERPs as well. Audiovisually congruent speech information was associated with reduced amplitude measures in early to late components in comparison to visually unspecific information. Age-related amplitude modulations were found, with enhanced amplitude measures in older adults in early processing stages and reduced amplitude measures in later stages. The occurrence of a deviant target location was associated with larger amplitudes in both early (P1-N1) and late (P2-N2) amplitude measures. In the following, specific results regarding audiovisual speech processing, dynamic switches in talker location, and age-related changes are discussed in detail.

### 4.1 Audiovisual speech comprehension in static and dynamic settings

Looking at the static setting, in which no switches in target location occurred, we observed a benefit of audiovisual information, which was being reflected both behaviorally and electrophysiologically. There were faster and more accurate responses with audiovisually congruent stimuli compared to visually unspecific or still face stimuli. In terms of accuracy, the benefit of congruent in contrast to unspecific information was especially visible in older, but not younger adults. On the one hand, this could be due to ceiling effects in the younger group, with error rates of less than 3%. On the other hand, it has been shown that younger adults are less biased by ambiguous visual information (Cienkowski & Carney, 2002), while older adults need high quality visual input in order to benefit from it (Tye-Murray et al., 2011). Generally, our study is in line with the notion that audiovisually congruent information supports effective speech comprehension by integrating coherent and redundant information from both modalities (Besle et al., 2004; Bronkhorst, 2015; Campbell, 2008; Lindström, 2012; van Wassenhove et al., 2005). In addition, Stekelenburg and Vroomen (2007) argued that enhancement due to successful integration does not primarily depend on the congruency of both modalities, but on the predictive value of the visual input. That is, additional visual input only leads to a decreased auditory threshold (Grant & Seitz, 2000) and enhanced intelligibility (Schwartz et al., 2004) when it allows for a valid prediction of the subsequent auditory information. Consistently, our visually unspecific condition demonstrates that preceding lip movements which might serve as a cue to the onset of the following speech sound, but which are unrelated to the auditory information, do not seem to be sufficient to result in behavioral benefits. In conclusion, not the timing, but the content is important for a facilitation in speech comprehension.

When presented within the dynamic setting, a reversion of the benefit of audiovisually congruent information became visible as soon as an unexpected change in target talker location occurred. Previous research in the auditory modality has already demonstrated that switches in target talker location deteriorate speech comprehension in cocktail-party situations (Best et al., 2008). This has been attributed to the re-analysis of the auditory scenary and the re-focussing of auditory attention after a switch has occurred (Getzmann et al., 2015; Koch et al., 2011; Lin & Carlile, 2015). Additionally, when presented with the audiovisually congruent stimuli in the present study, the fixated faces at the (central) standard location became temporarily incongruent in case of a location switch: Instead of the expected visual speech information (the target words “yes” or “no”), a deviant visual input occurred not matching the expected subsequent auditory input. Incongruent audiovisual input has been shown to result in increased response times compared to congruent stimuli (e.g., Altieri, Lentz, Townsend, & Wenger, 2016). This obvious disadvantage of audiovisual speech in dynamic multi-talker scenarios corresponds to results of a previous study, in which speech recognition in a multi-talker context with changing talkers was slower in an audio-visual condition compared to an audio-only condition (Heald & Nusbaum, 2014). The authors attributed this effect to an enhanced talker variability when presented with audiovisual stimuli Overall, our results suggest a behavioral benefit of audiovisually congruent information, when the target was presented from the expected location.

The deviance-related increase in response times with audiovisual information compared to still faces or unspecific lip movement was more pronounced in older than in younger adults, suggesting that they suffered more from the sudden switch in target talker location than the younger group. That is, the benefit shown by the older group in the static setting was reversed when a change of target talker occurred. In line with this, previous research has shown that older adults experience higher conflict costs in multi-talker scenarios, resulting in less accurate responses, when the attentional load is high (Passow et al., 2012), and higher response times following distraction due to unpredictable sound modulations (Volosin et al., 2017) or deviant target positions (Getzmann et al., 2015). It could be argued that these age-specific modulations are (at least partly) based on reduced basic unisensory and cognitive abilities, as also found in the present study in hearing and vision (but not lip reading) as well as in cognition. In fact, there are several studies demonstrating that audiovisual speech facilitation was independent of age, when unisensory performance was controlled for (Sommers et al., 2005; Tye-Murray et al., 2016). In another study with two groups of older adults with either hearing impairment or normal hearing, similar benefits from the audiovisual signal were found when controlling for unimodal deficits, that is, irrespective of hearing status (Tye-Murray, Sommers, & Spehar, 2007). Thus, it cannot be completely ruled out that basic unisensory and cognitive abilities did not also play a role in audiovisual speech comprehension in dynamic settings. However, in our study, we did not find correlations of unisensory and cognitive abilities and behavioral performance for the two age groups, implying that the observed effects of audiovisually congruent information should rather be associated with visual facilitation and not unisensory or cognitive differences.

### 4.2 ERPs to audiovisual speech onset

The above-mentioned behavioral patterns are reflected in the ERP measures. In the static setting, the benefit due to audiovisually congruent information in comparison to visually unspecific information was associated with a reduction in the early P1-N1 complex, but only in the older group. This observation is in line with past research and has been interpreted as an indicator of facilitated speech processing at an early perceptual level, where both modalities of speech information are integrated (Baart, 2016; Ganesh et al., 2014; Stekelenburg & Vroomen, 2007; van Wassenhove et al., 2005). Moreover, this early integration of both modalities could enable a more effective pre-attentive filtering of sensory information (e.g., Lebib et al., 2003). Observing this effect only in the older group is in line with a previous EEG study in which larger reductions in P1 amplitude in older compared to younger participants were found when being presented with audiovisual versus auditory-only speech (Winneke & Phillips, 2011). In the dynamic setting, amplitudes were enhanced for deviant compared to standard trials. Yet, the P1-N1 reduction associated with audiovisually congruent stimuli compared to visually unspecific information remained present for the older group.

This is in line with findings by Lebib and colleagues (2003), revealing larger P1 amplitudes for incongruent audiovisual stimuli when the input from both modalities was harder to distinguish. This was interpreted as an indicator for early detection of non-redundant audiovisual information, with the lip movement being processed even before vowel sound onset. According to the “analysis by synthesis” approach by Wassenhove and colleagues (2005) audiovisual integration is suggested to take place as soon as the N1, that is, on a perceptual level. Several studies could further show that binding processes can be observed at the time interval of the N1 peak, mainly being driven by the anticipation due to the preceding visual speech input (Ganesh et al., 2014; Stekelenburg & Vroomen, 2007).

Later in the processing cascade, the N1-P2 amplitude measures were smaller with visually unspecific compared to audiovisually congruent stimuli, but only in the dynamic setting. This is in line with previous studies, showing amplitude reductions in both the N1 and P2 amplitude when presenting an audiovisual stimulus, making the components an indicator for successful audiovisual integration (e.g., Baart, 2016; Baart et al., 2014; Ganesh et al., 2014; Klucharev et al., 2003). Taken together, the results from both P1-N1 and N1-P2 clearly demonstrated the difference between congruent, informationally redundant and unspecific, informationally irrelevant visual input in the context of audiovisual speech perception. Despite its temporal precedence relative to the auditory signal, unspecific visual information did not provide valid or helpful information, but rather seemed to demand more cognitive resources to process.

Finally, in the static and dynamic setting, a reduced P2-N2 amplitude was observed with audiovisually congruent relative to unspecific stimuli. One possible explanation may be that audiovisually congruent information results in a reduction of the required cognitive control for the processing of audio-visual language in a mixture of speech stimuli. However, independent of the presented visual input, we found an increase in P2-N2 amplitude in the dynamic setting when the target stimulus came from the deviant position. This suggests that a higher amount of cognitive control was needed when the target location changed. Similar results have been shown for non-speech auditory stimuli, where N2 deflections were elicited when presenting preceding visually incongruent information (Lindström, 2012). The effect may as well have been driven by modulation of the P2 component.

Finally, it should be noted that while most findings from previous studies focus on singular ERP components, we chose to analyze mean-amplitude-to-mean-amplitude ERP measures. We thereby accounted for a pronounced CNV-like preparatory activity prior to sound onset, which has also been addressed in previous studies on audio-visual speech processing (e.g., Teder-Salejarvi et al., 2002). Relatively strong high-pass filtering of 1 Hz or above has been applied to filter out effects of slow potentials like this (e.g., Ganesh et al., 2014; Winneke & Phillips, 2011). However, this filtering comes with the risk of introducing unwanted filter effects (the effect of different filters on ERPs in audiovisual speech experiments has previously been investigated by Huhn, Szirtes, Lorincz, & Csépe, 2009; for general thoughts on high-pass filtering see Luck, 2014). We therefore chose a different approach and analyzed amplitude to amplitude measures, which has also been done before to account for parietal buildup (Pilling, 2009) or baseline effects (Kokinous et al., 2015). Consequently, choosing this different measurement for analyses sacrifices temporal precision to a certain extent, but makes it possible to further investigate preparatory activity in speech processing.

### 4.3 Preparatory ERP activity preceding auditory speech onset

Our results showed stronger CNV amplitudes for audiovisually congruent information than for visually unspecific stimuli. This suggests more pronounced preparatory processes, when the visual information was known to be relevant. In this regard, it should be noted that we chose a blocked design in our experiment. Thus, participants would know that the lip movement contained valid information as opposed to the still faces and simple mouth opening and closing movements in the other two conditions. The CNV is typically elicited, when a preceding stimulus reliably predicts a subsequent stimulus, containing task-relevant information (Brunia, van Boxtel, & Böcker, 2011; Walter et al., 1964). Anticipatory slow-waves such as the CNV in the context of audiovisual integration have been sparsely studied so far, although their occurrence is known (Teder-Salejarvi et al., 2002). Our findings are in line with the notion of the CNV as an expectancy and anticipation-related component. In particular, the more pronounced CNV amplitude for audiovisually congruent stimuli might reflect a heightened attentional focusing towards the visual speech input and might also be an indication of successful integration as soon as the target sound would be presented.

In the dynamic setting, the enhancement of the CNV in audiovisually congruent compared to visually unspecific stimuli could only be observed for standard, but not for deviant trials. Naturally, the present experimental setup included a dynamic interplay between expectations about the upcoming speech content and the subsequent effects of (spatial) attention. In the dynamic setting, participants expected the target to be presented from a mostly constant position and the focusing of attention on this expected target position could have driven stronger audiovisual facilitation in such standard trials. In deviant trials, however, the identification of a mismatch between the expected visual speech content (i.e., lip movements associated with the target) and the observed visual speech was possible even before the auditory speech content occurs. The detection of an early visual mismatch at the expected target position might consequentially have triggered the shift of attention towards an alternative target location. Kononowicz and Penney (2016) suggestedd, that the CNV reflects a representation of timing processes that can also be observed in subsequent ERP components such as the N1 and P2. The preceding visual input occurring in natural audiovisual speech enables preparatory activity, that would be reflected by the CNV. The predictive value of preceding visual input in speech has been demonstrated in several studies (e.g. Pilling, 2009; Stekelenburg & Vroomen, 2007; van Wassenhove et al., 2005). As discussed by Ganesh and colleagues (2014), audiovisual speech processing undergoes several processing steps, involving attentional processes, the subsequent binding of two unisensory inputs and finally, the integration and congruent perception of audiovisual information.

Finally, we were not able to find clear age effects, although previous findings suggest that preparatory attentional processes in general decline with age, being reflected in diminished CNV amplitudes (for overview, see Wild-Wall, Hohnsbein, & Falkenstein, 2007), especially when a widened focus of attention was needed, that is, in a divided opposed to a selective attention task (Wild-Wall & Falkenstein, 2010). Accordingly, a reduced CNV of older participants has primarily been observed when attention had to be divided between talkers in a cocktail-party situation (Getzmann et al., 2016). In the present task, the well-pronounced CNV to audio-visually congruent speech suggest intact preparatory attention and gating of subsequent task-related auditory speech information in the older group of participants.

### 4.4 Age effects

We observed general age effects in both behavioral and neuronal measures, demonstrating well-assessed age-related decline in performance for older adults in both cocktail-party scenarios (Getzmann & Wascher, 2016) and (audiovisual) speech processing (Sekiyama et al., 2014; Winneke & Phillips, 2011). In P1-N1, or early stages of processing, we found larger amplitude measures for the older group with simultaneously decreased P2-N2 amplitude measures. The increase in P1 and decrease in N2 has repeatedly been associated with an age-related deficit of inhibitory processing of irrelevant information in line with the Inhibitory Deficit Hypothesis, suggesting an age-related deficit in inhibitory processing in early stages that are associated with sensory gating (Friedman, 2011; Getzmann et al., 2015; Stothart & Kazanina, 2016). Stothart and Kazanina (2016) suggested, that resolving audiovisual incongruency is intact, but demands higher temporal and neural costs in older adults. In the present study, we also observed a decrease in P1-N1 amplitude measure in the older, but not in the younger group. This is in accordance with other findings demonstrating stronger modulations of the P1 component in the older group (Winneke & Phillips, 2011). As discussed above, this decrease in amplitude can be interpreted as cross-modal sensory gating (Lebib et al., 2003). Importantly, while differences between both age group have been observed in P1-N1 and P2-N2 measures, we still found a decrease from early to late processing stages. The results indicated that especially in an early processing phase associated with sensory gating older adults rely on congruent visual information. Together with previous findings, our results suggest that audiovisually congruent information provides the possibility to compensate for age-related inhibitory deficits at an early processing stage. Opposed to research showing an age-related modulation of N1 an P2 amplitude (see Friedman, 2011), we could not assess a clear age-effect on N1-P2. Unfortunately, using mean-amplitude-to-mean-amplitude measures due to preparatory activity before sound onset (CNV) makes it difficult to assess specific modulations on the single ERPs. While P2 amplitudes tend to be enhanced due to age, N1 modulations have been ambiguous throughout studies (for review see Friedman, 2011). In fact, it may be possible that age-related enhancement or decrease in both amplitudes may have been cancelled out in the mean-amplitude-to-mean-amplitude analysis.

### 4.5 Conclusion

To summarize, the ERP results from our study indicate that congruent visual speech information is actively used in preparation for the subsequent auditory input (CNV), enables early audiovisual integration (P1-N1), and a facilitation of cognitive control processes in the further processing of multi-talker speech (P2-N2), leading to a better speech perception as reflected by the enhanced performance (i.e., accuracy and response times). However, these benefits are only visible as long as the multi-talker scenario remains stable. In contrast, unexpected switches in target talker location lead to an audiovisual incongruency, cancelling the benefits of successfully integrated congruent audiovisual speech. This was observable in amplitude modulations in the P1-N1, P2-N2, and CNV measures and ultimately also in deteriorated performances measures. Although we found age-related changes in the processing of the speech stimuli, probably due to inhibitory deficits and worsened resource allocation, we still found that older adults were able to profit from congruent audiovisual information. The benefits were visible especially in early phases of processing (P1-N1), indicating a more effective pre-attentive filtering of sensory information, when presented with relevant additional visual input.

## Supporting information

Supplementary Analysis

## Acknowledgements

The authors are grateful to Ludger Blanke, Christele Motcho and Nina Abich for preparing the software and parts of the electronic equipment and to Stefan Weber, Kristin Koberzin, Vera Birgel and Denise Böhle for their help in running the experiment. Further, they would like to thank Nina Abich, Michaela Djudjaj, Kimberly Freytag and Bianca Zickerick for allowing to use their faces and voices as stimulus material.

## DECLARATIONS

### Contributions

Stephan Getzmann, Alexandra Begau, Edmund Wascher and Daniel Schneider designed the study, Alexandra Begau collected the data and conducted the data analysis supported by the helpful recommendations and discussions by and with Laura Klatt. All authors intensively discussed the results and their interpretations, the manuscript was drafted by Alexandra Begau, and revised by all authors.

### Conflicts of interest/Competing interests

All authors disclose no actual or potential conflicts of interest including any financial, personal, or other relationships with other people or organizations that could inappropriately influence (bias) their work.

## Funding

This work was supported by a grant from the Deutsche Forschungsgemeinschaft (GE 1920/4-1).

## Notes

### Summary of Updates

Result section substantially revised, Supplementary analyses added, updated figures and tables updated discussion

