## Supplementary Analysis for "Do congruent lip movements facilitate speech processing in a dynamic audiovisual multi-talker scenario? An ERP study with older and younger adults"

Supplementary data

**Supplementary analysis 1: Correlations of unisensory and cognitive abilities and behavioral measures**

In order to test whether behavioral performance in the experiment was related to unisensory abilities and basic neuropsychological measures of participants, Spearman-Brown correlations (Spearman & Spearman, 1904) were calcuated. P-values were FDR corrected (Benjamini & Hochberg, 1995). For analyses, data were collapsed across the conditions of visual information, settings, and with regards to the dynamic setting, across target positions. The results (table S1) did not reveal significant correlations (all $p_{adj} > .05$).

**Supplementary analysis 2: Comparing standard trials in static vs. dynamic setting**

In order to assess whether the setting had an influence on the processing of stimuli, we compared standard trials in the static and dynamic setting for the behavioral measures (in a 2 (age) × 3 (visual information) × 2 (setting) ANOVA) and for the ERPs (in a 2 (age) × 2 (visual information) × 2 (setting) ANOVA; here, we excluded still face stimuli), with focus on effects concerning the experimental setting. For response times, the analysis revealed higher response times in the dynamic setting, $M$ = 1724.83 ms, $SE$ = 18.40, compared to the static setting, $M$ = 1672.54 ms, $SE$ = 15.21 (table S2). No effects of setting were found for accuracy. Looking at ERP measures, the setting did not influence P1-N1 and N1-P2 measures. For the P2-N2 measure, we found an interaction of age with the setting. In the younger group, there was no difference between both settings, $t(65) = -1.88$, $p_{adj} = .065$, $g = 0.23$, while in the older group, amplitudes were larger in the dynamic compared to the static setting, $t(59) = 2.35$, $p_{adj} = .044$, $g = 0.30$.

To summarize, we only found a significant interaction of setting for the P2-N2 measure with follow-up analyses revealing larger amplitude measures in the dynamic compared to the static setting only within the older group. This could be explained by a higher task demand due to the dynamic setting that was only present for the older participants.

**Supplementary analysis 3: ERP latencies.**

The present data analysis was focused on amplitude modulations of the P1, N1, P2 and N2 components, but there is also some evidence of latency modulations of said ERP measures in audiovisual speech perception (e.g. Baart, 2016; Lindström, 2012; Winneke & Phillips, 2011). Therefore, an additional analysis of ERP latencies was conducted. In pre-specified time-windows, the latencies of the maximum or minimum peaks, respectively, were estimated for each subject in each condition. The time-windows were 20 to 120 ms for P1, 75 to 175 ms for N1, 150 to 250 ms for P2 and 250 to 350 ms for N2. The statistical analysis was identical to the one described for behavioral and amplitude measures in section 2.7.2.

For the *static setting*, we conducted a 2 (age) × 2 (visual information) ANOVA for each component (figure S3; table S3). For the P1, no latency differences were found. For the N1, the analysis revealed an interaction of visual information and age group. However, pairwise comparisons were not significant for neither the older, $t(19) = -2.25$, $p_{adj} = .073$, $g = -0.48$, nor the younger group, $t(21) = 1.33$, $p_{adj} = .197$, $g = 0.27$. For the P2, larger latencies were found in the older, $M_{old} = 203.80$ ms, $SE = 3.16$, compared to the younger group, $M_{young} = 179.27$ ms, $SE = 2.31$. Also for the N2, larger latencies occurred in the older, $M_{old} = 317.00$ ms, $SE = 4.88$, compared to the younger group, $M_{young} = 283.95$ ms, $SE = 4.82$.

For the *dynamic setting*, we conducted a 2 (age) × 2 (visual information) × 2 (target position) ANOVA for each component (figure S4; table S4). For P1, no significant differences between the conditions were found. For N1, the analysis revealed an interaction of visual information with the age group. Pairwise comparisons showed that while in the older group, larger latencies occurred for stimuli with audiovisually congruent, $M_{congr} = 102.30$ ms, $SE_{congr} = 2.91$, compared to visually unspecific information, $M_{unspec} = 108.80$ ms, $SE_{unspec} = 2.79$, $t(39) = -2.69$, $p_{adj} = .021$, $g = -0.42$, we observed a reversed pattern in the younger group, $M_{congr} = 109.00$ ms, $SE_{congr} = 3.39$, $M_{unspec} = 102.77$ ms, $SE_{unspec} = 2.73$, $t(43) = 2.02$, $p_{adj} = .049$, $g = 0.30$. For P2, the analysis revealed later amplitudes in the older, $M_{old} = 201.38$ ms, $SE = 2.54$, compared to the younger group, $M_{young} = 184.30$ ms, $SE = 1.69$. The N2 peaked later in the older, $M_{old} = 316.88$ ms, $SE = 3.24$, compared to younger group as well, $M_{young} = 295.18$ ms, $SE = 3.47$.

In sum, in both settings, the earliest latency modulation became present at the N1 component, with an interaction of visual information and the age group. Larger peak latencies were generally present in older adults for the later components P2 and N2 (which is in line with the literature, e.g. Anderer, Semlitsch, & Saletu, 1996). No additional effects of target position were found.

SUPPLEMENTARY TABLES

Table S1

*Correlations between unisensory and cognitive abilities and performance measures in the older and younger group*

| | Older group | | Younger group | |
|---|---|---|---|---|
| Variables | Response time | Accuracy | Response Time | Accuracy |
| | *ρ* | *ρ* | *ρ* | *ρ* |
| Visual Acuity[1] | .18 | .24 | -.01 | .01 |
| Hearing level[2] | .14 | -.05 | -.34 | .06 |
| MoCA Score[3] | -.09 | .29 | -.23 | -.02 |
| Digit repetition forward[4] | -.05 | .09 | -.48 | .18 |
| Digit repetition backward[4] | -.13 | .11 | -.26 | .28 |

*Notes.* Spearman ρ correlation coefficients for correlation of unisensory and cognitive abilities and performance measures, that is response time and accuracy. Data were collapsed over all within-factors.

[1] *Landolt C optotypes* in 1.5 m distance, binocular.

[2] Pure-tone audiometry (Oscilla USB 330; Inmedico, Lystrup, Denmark), mean across both ears for frequencies 125 to 4000 Hz.

[3] *Montreal Cognitive Assessment* (Nasreddine et al., 2005)

[4] Digit Span subtest from the German Adaptation of the *Wechsler Adult Intelligence Scale*; Hamburg-Wechsler Intelligenztest für Erwachsene – Revision 1991; HAWIE-R, Tewes, & Wechsler, 1991.

all $p > .05$ (after FDR correction).

Table S2

*ANOVA including standard trials in the dynamic and static setting.*

| | Accuracy | Response times | | P1-N1 | | N1-P2 | | P2-N2 | |
|---|---|---|---|---|---|---|---|---|---|
| conditions | *F* | *F* | $\eta_p^2$ | *F* | $\eta_p^2$ | *F* | $\eta_p^2$ | *F* | $\eta_p^2$ |
| age | 5.85 | **9.43**** | .19 | 3.83 | .09 | **4.62*** | .10 | **8.62**** | .18 |
| visual information | **13.54** | **79.01***** | .66 | **19.89***** | .33 | 2.23 | .05 | **44.68***** | .53 |
| setting | 0.04 | **14.39***** | .26 | 0.04 | <.01 | 0.87 | .02 | 0.20 | <.01 |
| age × visual information | 6.54 | **5.97**** | .13 | **6.15**** | .13 | 0.07 | <.01 | 0.85 | .02 |
| age × setting | 0.001 | 0.95 | .02 | 1.64 | .04 | 0.01 | <.01 | **8.83*** | .18 |
| Visual information × setting | 0.32 | 0.60 | .01 | 1.50 | .04 | <0.01 | <.01 | 1.23 | .03 |
| age × visual information × setting | 1.46 | 0.69 | .02 | 1.26 | .03 | 0.02 | <.01 | 0.96 | .02 |

*Notes.* 2 (young vs. old) × 2 (congruent vs. unspecific visual information) × 2 (static vs. dynamic setting) repeated-measures ANOVA. In the dynamic setting, only standard trials were used for the analyses. For accuracy and response time, the additional condition *still faces* was included for the factor of visual information. Degrees of freedom for analyses with ERP components was 1,40 for all factors. Degrees of freedom for behavioral analyses: age, setting and age × setting: 1,40 (response times), 1.00 (accuracy, except 1,40 for age); visual information and age × visual information: 2,80 (response times), 1.87 (accuracy); visual information × setting and age × visual information × setting 1,80 (response times), 1.89 (accuracy). Note, that a nonparametric ANOVA was calculated for accuracy, since the normality assumption was violated. Here denominator degrees of freedom are set to infinite. For response times, Greenhouse-Geisser correction was applied to age × setting, age × visual information × setting. Reporting uncorrected degrees of freedom.

* $p < .05$, ** $p < .01$, *** $p < .001$.

Table S3.

*ERP latency analyses for the static setting.*

| conditions | P1[1] | N1[1] | P2[2] | | N2[1] |
|---|---|---|---|---|---|
| | *F* | *F* | *F* | $\eta_p^2$ | *F* |
| age | 0.44 | 0.20 | **24.61*** ** | .38 | **16.46*** ** |
| visual information | 3.78 | 0.48 | 0.90 | .02 | 1.28 |
| age × visual information | 0.43 | **7.47** ** | 2.41 | .06 | 2.13 |

*Note.* 2 × 2 ANOVA, with age (young vs. old) and visual information (audiovisually congruent vs. visually unspecific) in the static setting.

[1] A nonparametric ANOVA-type statistics were calculated for P1, N1 and N2 due to violation of normality assumption, denominator degrees of freedom are infinite, thus reporting only nominator degrees of freedom, *df* = 1.00.

[2] *df* = 1,40 for all factors, except age: 1,39.96 (P1), 1,39.36 (N1), 1,39.99(N2).

** $p < .01$, *** $p < .001$.

Table S4.

*ERP latency analysis for the dynamic setting.*

| | P1 | N1 | P2 | N2 |
|---|---|---|---|---|
| | *F* | *F* | *F* | *F* |
| age | 0.04 | 0.01 | **17.03*** ** | **13.37*** ** |
| visual information | 0.82 | 0.07 | 0.27 | 1.05 |
| target position | 0.52 | 0.02 | 0.01 | 2.99 |
| age × visual information | 1.83 | **8.53** ** | 0.05 | 0.03 |
| target position × visual information | 0.46 | 0.16 | 0.21 | 1.10 |
| age × target position | 0.96 | 0.04 | 2.14 | 0.32 |
| age × visual information × target position | 0.42 | 0.92 | 0.33 | 0.27 |

*Note.* 2 (young vs. old) × 2 (congruent vs. unspecific visual information) × 2 (standard vs. deviant target location) repeated-measures ANOVA in the dynamic setting. Nonparametric ANOVA-type statistic, except for whole-plots factors, denominator degrees of freedom are infinite, thus reporting only nominator degrees of freedom.

Degrees of freedom: *df* = 1.00 for all factors, except age: 1,38.00 (P1), 1,39.84 (N1), 1,38.67 (P2), 1,37.26 (N2).

** $p < .01$, *** $p < .001$.

SUPPLEMENTARY FIGURES

**Figure S1.** Waveforms are shown for separately for the younger and older group, for audiovisually congruent and unspecific information in the static setting. The time interval shows length of one epoch, baseline interval was -1600 to -1500, i.e. prior to stimulus onset, sound onset was at 0 ms. The grey area marks the analyzed part of the ERP wave, that is, the CNV.

**Figure S2.** Waveforms are shown for separately for the younger and older group, for audiovisually congruent and unspecific information, and for standard and deviant trials in the dynamic setting. The time interval shows length of one epoch, baseline interval was -1600 to -1500, i.e. prior to stimulus onset, sound onset was at 0 ms. The grey area marks the analyzed part of the ERP wave, that is, the CNV.

**Figure S3.** ERP latencies in static setting: Peak latencies for P1, N1, P2 and N2 shown separately for the younger and older group, for audiovisually congruent and still visual information. Box plot whiskers mark 1.5 IQR, mean is indicated by the cross. Significant results are marked with vertical lines, *** $p < .001$.

**Figure S4.** ERP amplitudes in dynamic setting: Peak latencies for P1, N1, P2 and N2 shown separately for the younger and older group, for audiovisually congruent and still visual information, and for standard and deviant trials. Box plot whiskers mark 1.5 IQR, mean is indicated by the cross. Significant results are marked with vertical lines, * $p < .05$, *** $p < .001$.

Figure S1

*ERP waveforms with 0.1 Hz high-pass filter at electrode cluster Cz, FCz, Fz, FC1 and FC2 in static setting*

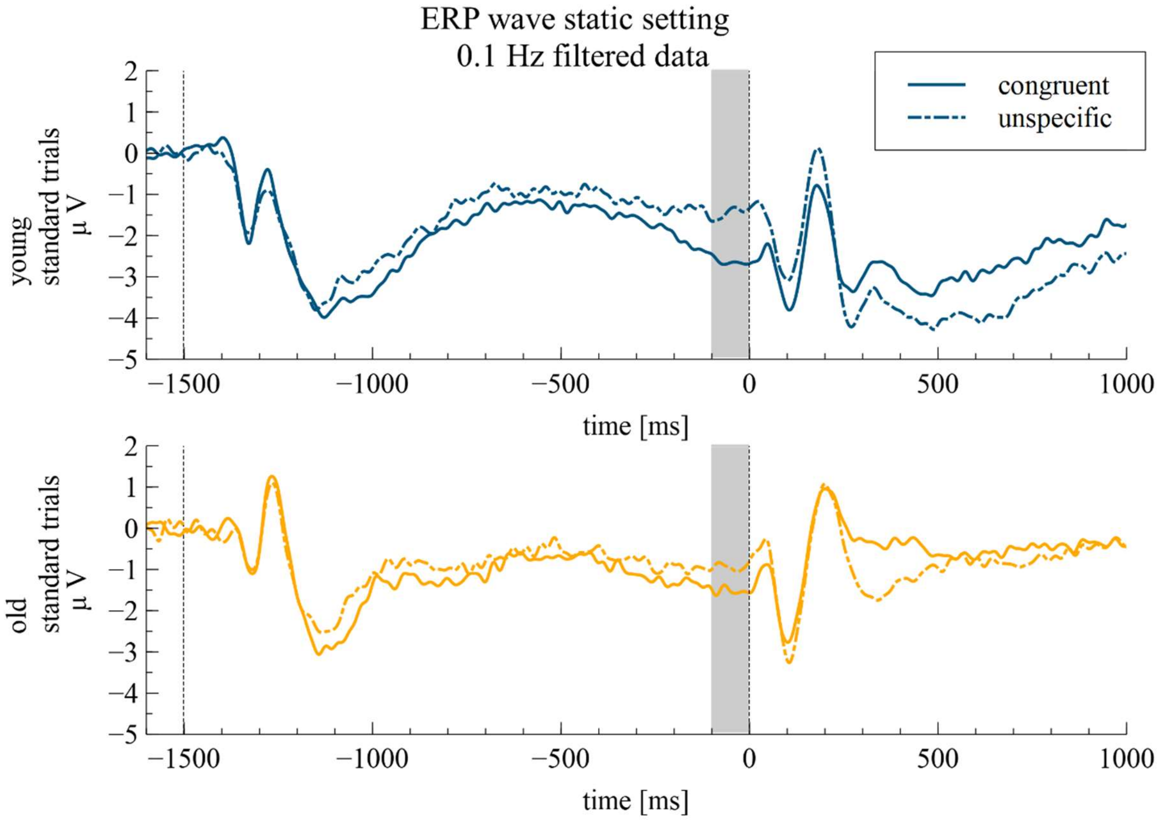

Figure S2

*ERP waveforms with 0.1 Hz high-pass filter at electrode cluster Cz, FCz, Fz, FC1 and FC2 in dynamic setting*

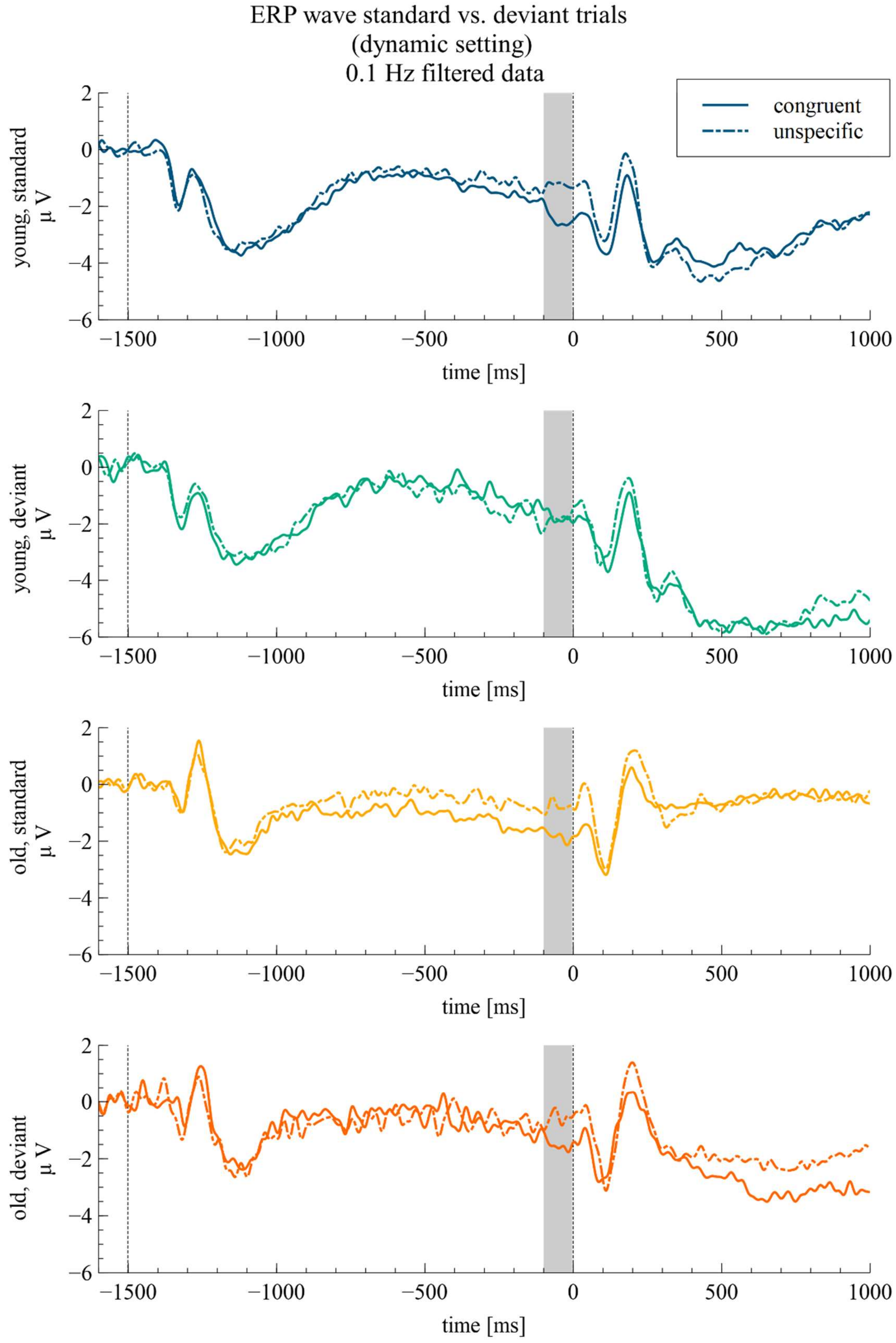

Figure S3

*ERP latencies in the static setting*

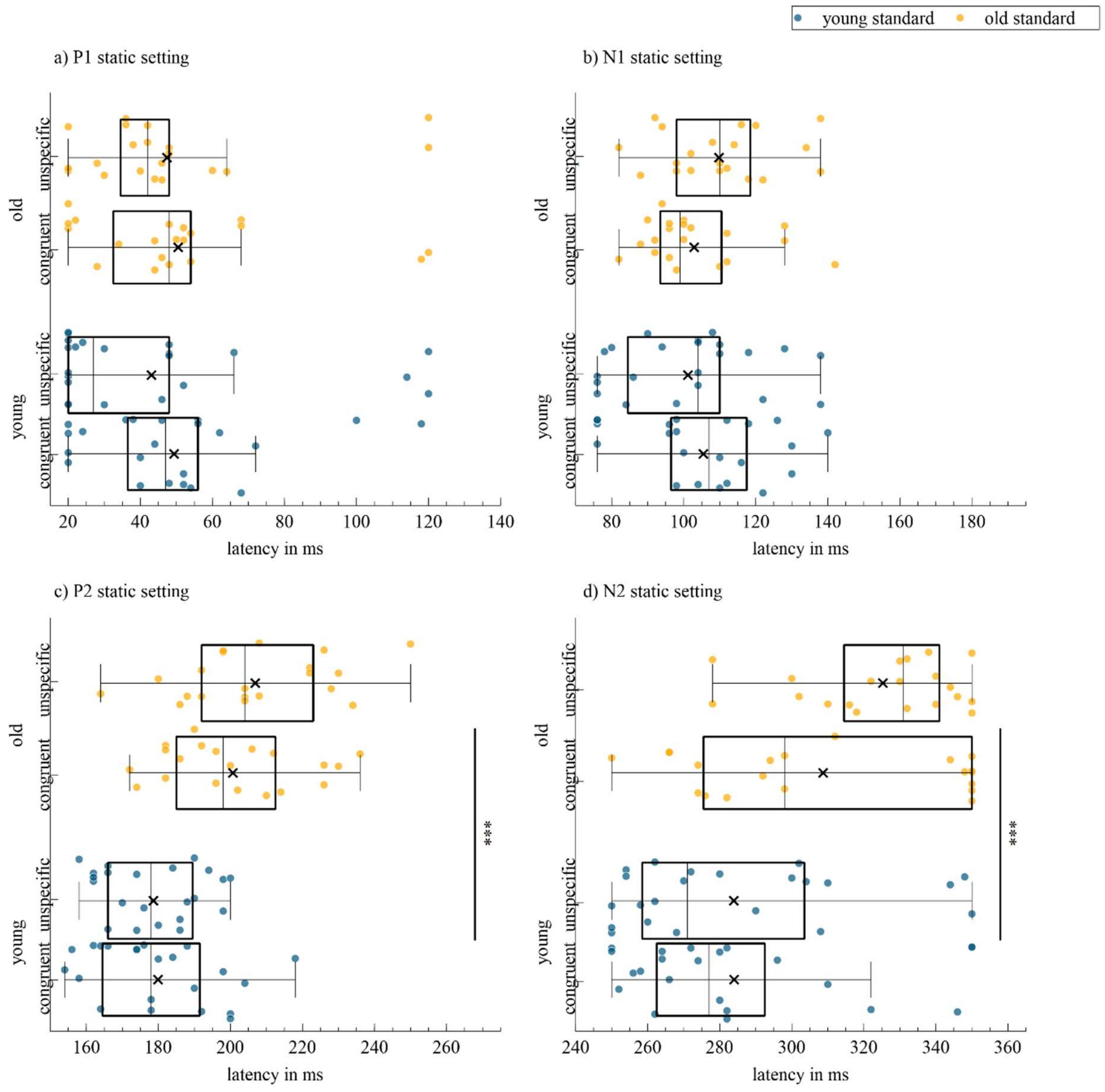

Figure S4

*ERP latencies in the dynamic setting*

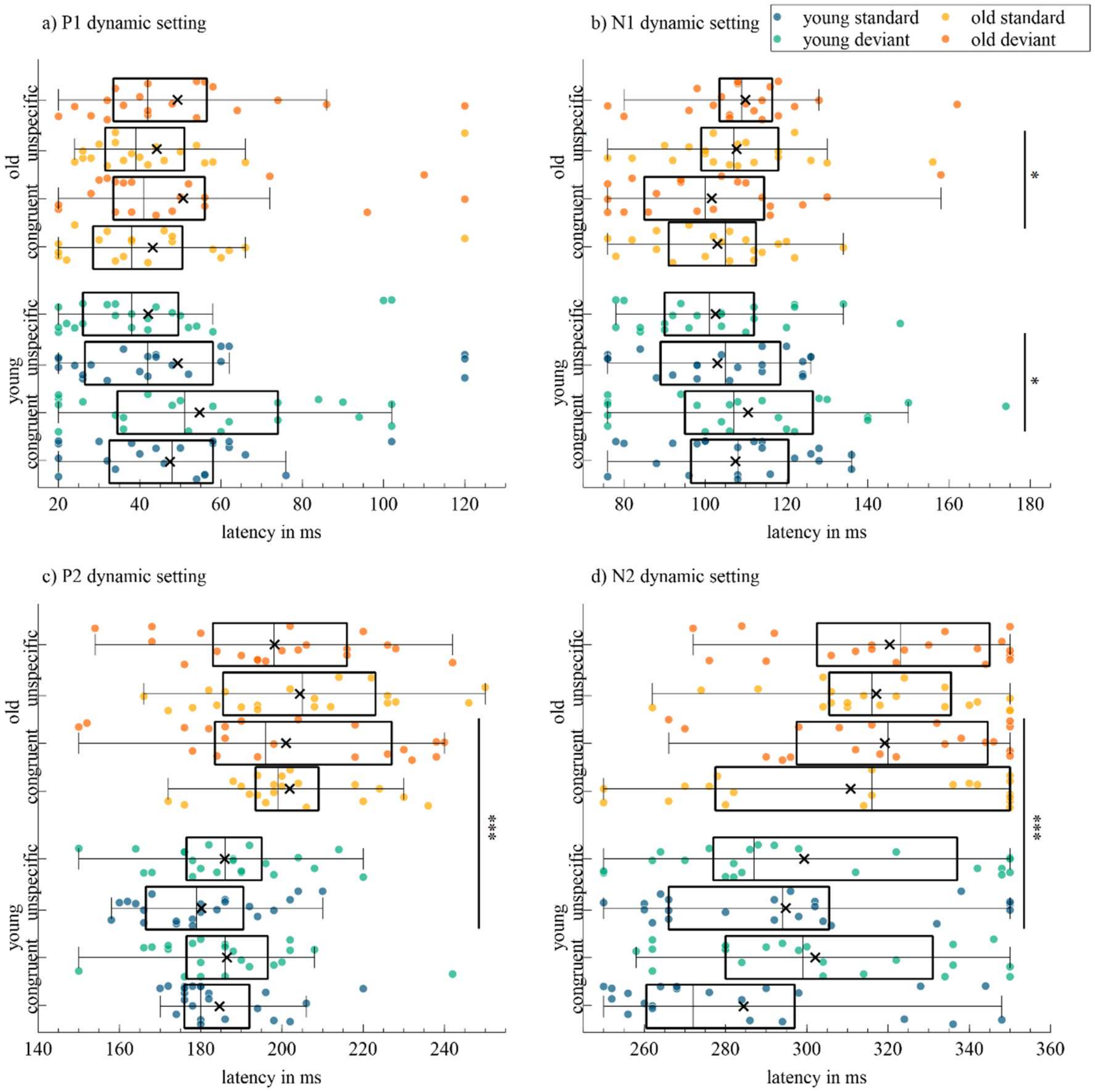
